# IRE1α and IRE1β Protect Intestinal Epithelium and Suppress Colorectal Tumorigenesis through Distinct Mechanisms

**DOI:** 10.1101/2025.05.01.651751

**Authors:** Ruishu Deng, Miao Wang, Thanyarat Promlek, Clementine Druelle-Cedano, Rabi Murad, Nicholas O. Davidson, Randal J. Kaufman

## Abstract

Intestinal epithelial cells (IECs) uniquely express two IRE1 paralogues, IRE1α and IRE1β, whose roles in intestinal physiology are incompletely understood. We examined the individual and cooperative functions of IRE1α and IRE1β in IECs using mice using intestine-specific deletion of *Ire1α* or germline *Ire1β* deletion, and subsequently with double *deleted Ire1α, Ire1β* mice. At baseline, intestine-specific *Ire1α* deleted mice and mice with germline *Ire1β* deletion exhibited no morphologic changes in small intestine or colon, but double deleted *Ire1α^-/-^Ire1β^-/-^* mice developed progressive intestinal and colonic injury and tumorigenesis. In contrast to single-deleted IECs, RNA-Seq from *Ire1α^-/-^Ire1β^-/-^* IECs revealed decreased expression of defense-associated mRNAs, together with increased expression of inflammatory and pathogenic mRNAs. Utilizing orthogonal models of intestinal tumorigenesis, reflecting either inflammatory-mutagenic injury (AOM-DSS) or spontaneous polyposis (APC^min^), we observed that loss of either intestinal epithelial *Ire1α* or of *Ire1β* alone produced a growth advantage, increasing tumor burden. IRE1α mediated splicing of *Xbp1* mRNA was maintained following *Ire1β* deletion but not in double deleted *Ire1α^-/-^Ire1β^-/-^* mice. Increased expression of either *Ire1α* or *Ire1β* mRNA was associated with improved survival in patients with colorectal cancer. Taken together our findings suggest IRE1 paralogues utilize essential but distinct mechanisms to safeguard intestinal homeostasis and suppress tumorigenesis.

## Introduction

Intestinal epithelial cells (IECs) rapidly self-renew and maintain high levels of protein secretory activity(^1^), relying heavily on an effective unfolded protein response (UPR) to ensure homeostasis (2). Inositol-requiring enzyme 1 (IRE1) is the most conserved UPR signaling arm (3), which in mammals involves two paralogues, IRE1α (*ERN1*) and IRE1β (*ERN2*) (4, 5). IRE1α is ubiquitously expressed, and its endoribonuclease (RNase) activity removes a 26 base unconventional intron from *Xbp1* mRNA, to produce a spliced mRNA *(Xbp1s)*, that activates the transcriptional activity of XBP1 to induce target genes (5–9). In contrast, IRE1β expression is restricted to epithelial cells in the intestine and lung (10, 11). However, the cellular distribution and independent versus cooperative functions of IRE1α and IRE1β in IECs remain incomplete.

Studies of the downstream targets of IRE1 revealed that IEC-specific *Xbp1* deleted mice display spontaneous enteritis and exacerbated DSS-induced colitis(12). Subsequently, increased intestinal tumorigenesis was observed in IEC-specific *Xbp1* deficient mice(13). However, hyperactivation of IRE1α was also observed in these mice and was considered a driver of both intestinal inflammation and tumorigenesis (14). Other studies showed that IEC-specific deletion of IRE1α exon 2 in mice was associated with spontaneous colitis and increased mortality (15) suggesting a protective physiological role for IRE1α-XBP1s in the intestinal tract. However, depleting IRE1α suppressed colon cancer in those IRE1α exon 2-deleted mice, indicating a pathogenic role in tumorigenesis(16). Although reduced tumor formation was observed in the colons of IEC-specific IRE1α exon 2-deleted mice in the AOM-DSS model (16), those mice developed spontaneous colitis and increased sensitivity to DSS-induced colitis (15). Consequently, the reduced tumor formation observed in IRE1α exon 2-deleted mice(16) is seemingly inconsistent with the increased intestinal tumor formation observed in IEC-specific *Xbp1*-deleted mice(13). These conflicting results raise questions as to whether and how IRE1α modifies colorectal cancer susceptibility.

IECs express IRE1β, in addition to IRE1α(4, 10). IRE1β was shown to protect mice from DSS-induced colitis (10), with roles in goblet cell development (17), efficient MUC2 folding and secretion (18), as well as the degradation of microsomal triglyceride transfer protein (MTTP) mRNA to maintain intestinal lipid homeostasis (19). However, a role for IRE1β in colorectal cancer formation has yet to emerge. In addition, it is unclear if IRE1β shares the same function as IRE1α in splicing *Xbp1* mRNA to produce *Xbp1s* (12, 13). IRE1β was shown to splice *Xbp1* mRNA as evidenced by reduced levels of *Xbp1s* mRNA in the colon of *Ire1β*^-/-^ compared to WT mice (17) but it was unclear if downstream target gene expression was reduced in those *Ire1β*^-/-^ mice. These observations raise questions regarding the physiological function IRE1β in relation to Xbp1 mRNA splicing and alternative unconventional splicing targets, including its role in maintaining intestinal homeostasis and tumor formation, both alone and in combinations with IRE1α.

To investigate these questions, we created doubly deleted, intestine-specific *Ire1α^-/-^Ire1β^-/-^* mice, which exhibit spontaneous colonic injury. We further found that experimental CRC was increased in both intestine-specific *Ire1α^-/-^* or germline *Ire1β^-/-^* singly-deleted mice. Additionally, unlike Ire1α, Ire1β did not splice *Xbp1* mRNA; rather, Ire1β suppressed mTORC1 signaling. Analysis of patient datasets indicated that higher levels of IRE1α or IRE1β expression was associated with improved survival in CRC patients. Overall, our findings suggest that IRE1α and IRE1β utilize essential distinct mechanisms to protect intestinal homeostasis and prevent tumorigenesis.

## Results

### *Ire1α* and *Ire1β* are simultaneously expressed in cell populations that regulate intestinal homeostasis

We utilized fluorescence *in situ* hybridization (FISH) to visualize the distribution of *Olfm4* mRNA, along with *Ire1α* and *Ire1β* mRNA and used Ki67, Lysozyme (LYZ), and MUC2 antibodies as markers for transit amplifying cells, Paneth cells, and goblet cells, respectively. We observed that *Ire1α* exhibited diffuse expression in villus IECs, with selective expression of *Ire1β* in specific cell populations (Fig.1&2). We further demonstrated *Ire1α* and *Ire1β* were coexpressed in goblet cells (Fig. 1A Fig. S1A), Paneth cells (Fig. 1B), transit amplifying cells (Fig. 2A), and intestinal stem cells (ISCs) (Fig. 2B). These IEC populations play pivotal roles in intestinal crypt development and intestinal repair, suggesting that IRE1α and IRE1β may each play key roles in maintaining intestinal homeostasis.

**Fig. 1.**
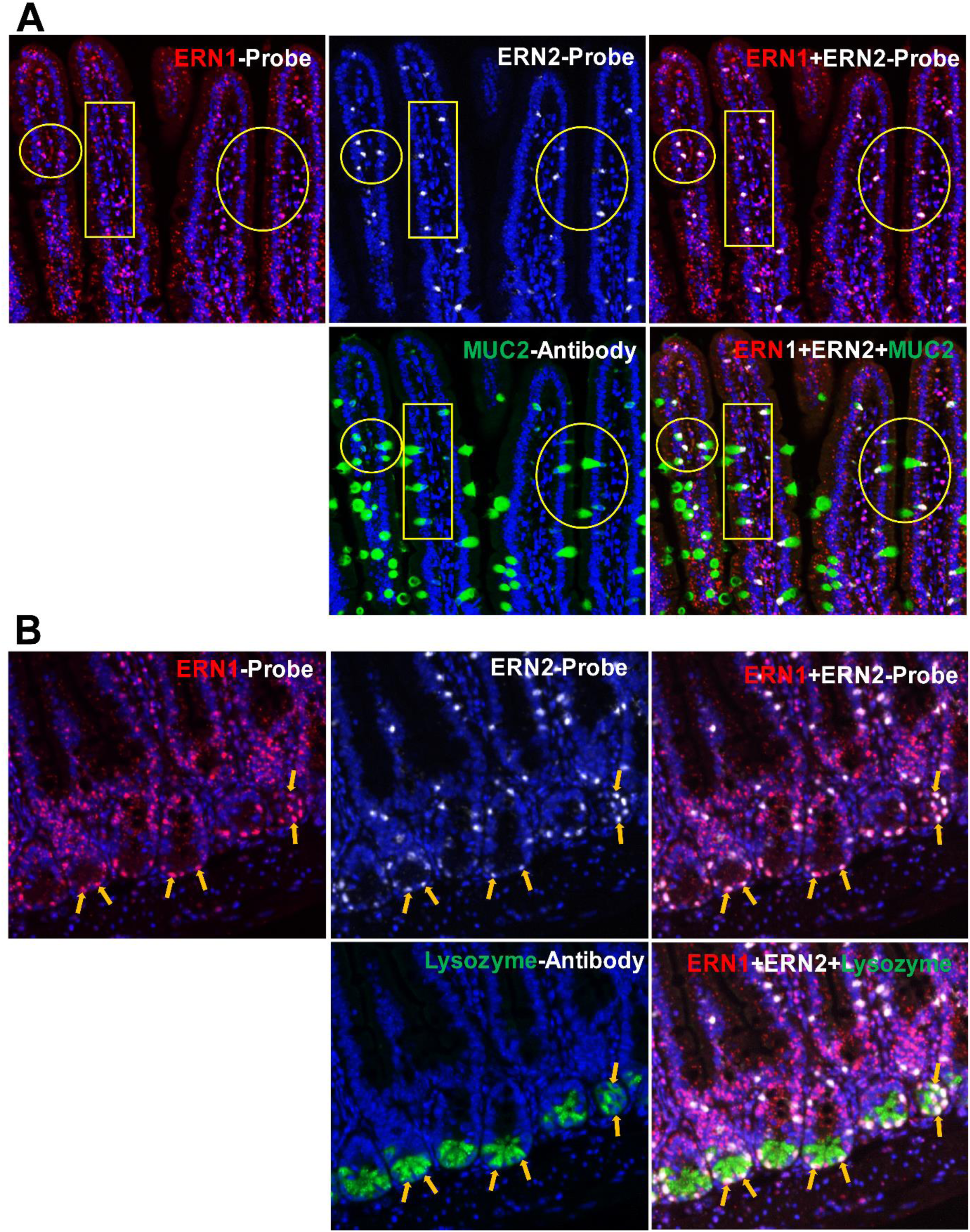
*Ire1α* and *Ire1β are co-expressed* in goblet cells and Paneth cells. The small intestines of 12-week-old mice were used for Fluorescent In Situ Hybridization to visualize the distribution of *Ire1α* and *Ire1β*. mRNA probes for *Ire1α* and *Ire1β* were applied to the same slide, along with antibody for MUC2 (A), LYZ (B), respectively, to identify the co-expression of *Ire1α* and *Ire1β* in goblet cells (A), Paneth Cells (B). The image represented five mice with an identical expression pattern of these two molecules.

**Fig. 2.**
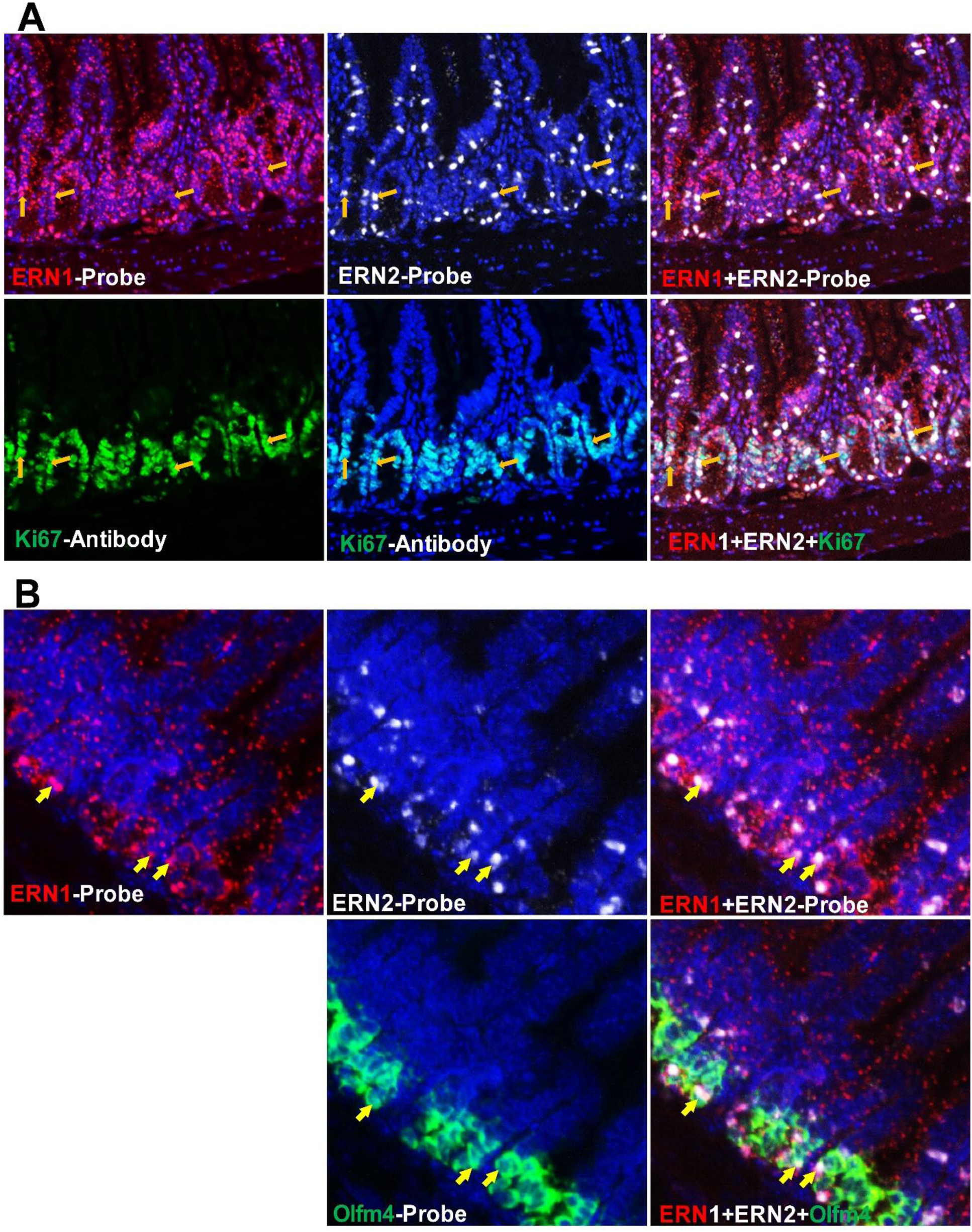
Co-expression of *Ire1α* and *Ire1β* occurs in transit amplifying cells and ISCs. The small intestines of 12-week-old mice were used for Fluorescent In Situ Hybridization to visualize the distribution of *Ire1α* and *Ire1β*. mRNA probes for *Ire1α* and *Ire1β* were applied to the same slide, along with antibody for Ki67 (A) to identify the co-expression of *Ire1α* and *Ire1β* in transit amplifying cells. *Olfm4* probe (B) was applied on the same slide as the *Ire1α* and *Ire1β* probes to confirm the expression of these two molecules in ISCs. The image represents five mice with an identical expression pattern of these two molecules.

### Intestine-specific deletion of *Ire1α* and *Ire1β* induces morphologic damage and alters gene expression in the small intestine

In order to determine the individual and cooperative roles of *Ire1α* and *Ire1β,* we generated intestine-specific *Ire1α^-/-^* mice and bred them with germline *Ire1β^-/-^* mice to generate IECs with intestine-specific deletion of both *Ire1α* and *Ire1β* (*Ire1α^-/-^Ire1β^-/-^*). Genome browser view of RNA-Seq genomic coverage confirmed successful deletion of *Ire1α* exons 16 and 17 and *Ire1β* exons 12 and 13 in IECs from *Ire1α^-/-^*, *Ire1β^-/-^* and *Ire1α^-/-^Ire1β^-/-^* mice (Fig. S1B). Body weights of chow-fed *Ire1α^-/-^* or *Ire1β^-/-^* mice were indistinguishable from wild type young and old mice (Fig. 3A; Fig. S2A). In contrast, body weights of double deleted *Ire1α^-/-^Ire1β^-/-^* mice were significantly decreased in aged mice (>40 wks old) but not in young mice (10-14 wks old) (Fig. 3A; Fig. S2A). Epithelial barrier function was compromised in both young and old *Ire1α^-/-^Ire1β^-/-^* mice as evidenced by increased serum FITC, but not in single deleted in *Ire1α^-/-^* or *Ire1β^-/-^* mice (Fig. 3B; Fig. S2B). No distinguishable morphological changes or spontaneous enteritis were observed in either *Ire1α^-/-^* or *Ire1β^-/-^* mice (Fig. 2C) or in young double deleted mice (Fig. S2C). However, small intestinal villi and crypt architectures were grossly disrupted in 100% of aged *Ire1α^-/-^Ire1β^-/-^* mice (Fig. 3C) and spontaneous enteritis was also observed in aged *Ire1α^-/-^Ire1β^-/-^* mice, a feature that was not observed in single deleted *Ire1α^-/-^* or *Ire1β^-/-^* mice (Fig. 3C). Analysis of the major small intestinal cell populations revealed that both Alcian blue positive goblet cells and LYZ positive Paneth cells were reduced in *Ire1α^-/-^Ire1β^-/-^* mice but were preserved in single deleted *Ire1α^-/-^* or *Ire1β^-/-^* mice. (Fig. 3D, E; Fig. S2D, E). In contrast, Ki67 positive transit amplifying cells were unchanged in single deleted *Ire1α^-/-^* or *Ire1β^-/-^* mice but increased in double deleted *Ire1α^-/-^Ire1β^-/-^* mice (Fig. 3F; Fig. S2F). These data indicate that combined intestine-specific deletion of *Ire1α* and *Ire1β* disrupts adult intestinal homeostasis with aging, while deletion of either *Ire1α^-/-^* or *Ire1β^-/-^* alone does not. These data suggest that IRE1α and IRE1β provide complementary functions to maintain intestinal homeostasis and barrier function.

**Fig. 3.**
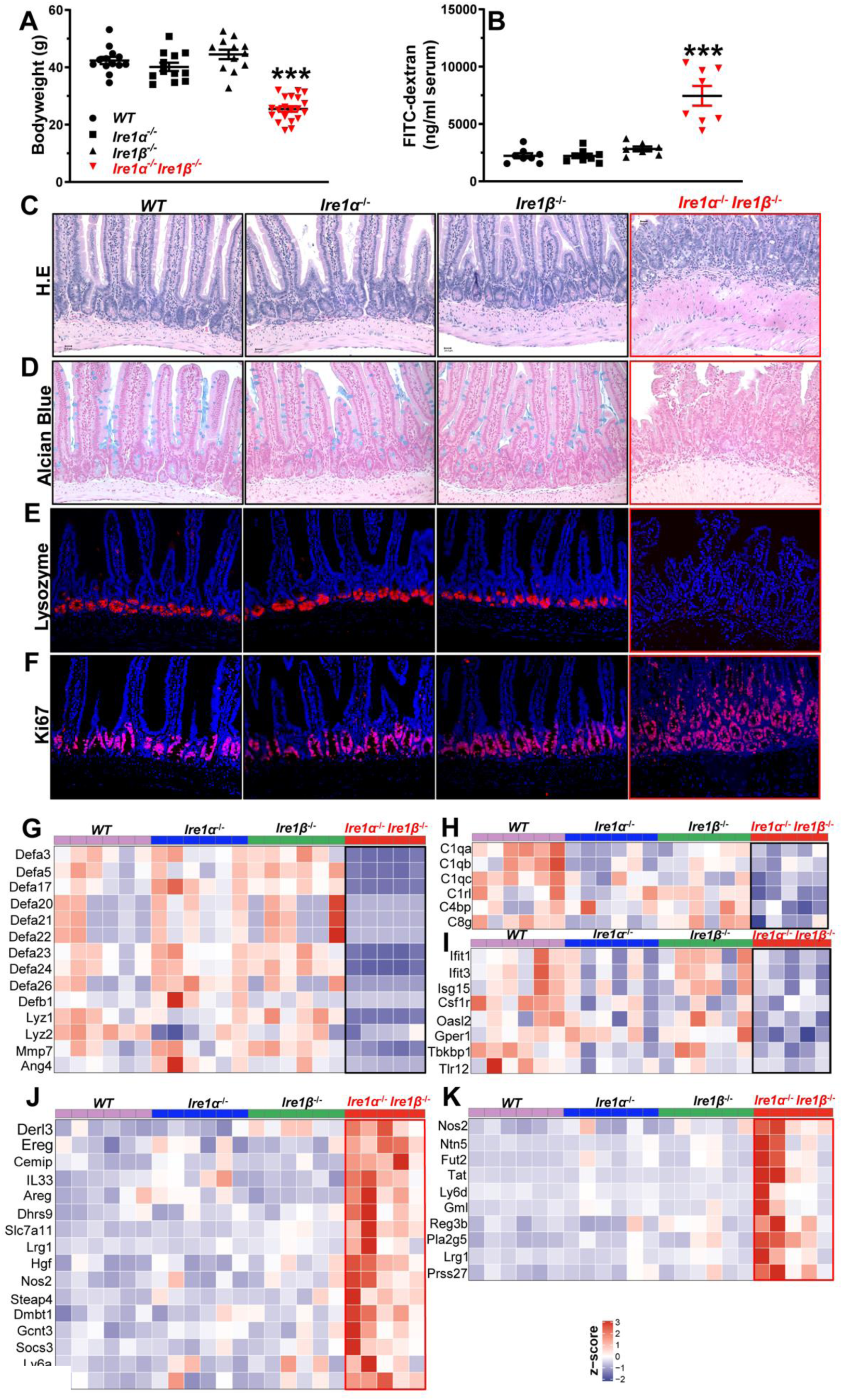
*Ire1α^-/-^ Ireβ^-/-^* mice display severe damage of small intestines and altered gene expression. (A) Body weight of aged (≥40 wks old) mice, WT n=13 mice, *Ire1α^-/-^* n=12 mice, *Ireβ^-/-^* n=12 mice, and *Ire1α^-/-^Ireβ^-/-^* n=20 mice is shown. Data shown are means ± SE. *** P < 0.0001 (B) Intestinal permeability was measured by serum FITC-dextran levels of mice (≥40 wks old), *WT* n=8 mice, *Ire1α^-/-^* n=8 mice, *Ireβ^-/-^* n=7 mice, *Ire1α^-/-^Ireβ^-/-^* n=8 mice. Data shown are means ± SE. *** P < 0.0001 (C) H&E staining shows architecture distortion in aged *Ire1α^-/-^Ireβ^-/-^* mice but not in age matched *WT, Ire1α^-/-^* or *Ireβ^-/-^* mice. Alcian blue staining (D) and immunofluorescence of LYZ (E) show absence of Alcian blue positive goblet cells and LYZ positive Paneth cells in aged *Ire1α^-/-^ Ireβ^-/-^* mice but not in age matched *WT*, *Ire1α^-/-^* or *Ireβ^-/-^* mice. (F) Immunofluorescence shows increased Ki67 positive cells in aged *Ire1α^-/-^Ireβ^-/-^* mice but not in *Ire1α^-/-^* or *Ireβ^-/-^* mice. The images shown are typical of 8 mice in each group. (G) Heatmap of selected genes depicts decreased defensins, LYZs, MMP7 and Ang4 in *Ire1α^-/-^Ireβ^-/-^* IECs. (H) Heatmap of selected genes shows decreased expression of indicated complements in *Ire1α^-/-^Ireβ^-/-^* IECs. (I) Heatmap of selected genes demonstrates decreased expression of innate immune response genes in *Ire1α^-/-^Ireβ^-/-^* IECs. (J) Heatmap of selected genes shows increased expression of genes involved in enteritis/colitis in *Ire1α^-/-^Ireβ^-/-^* IECs. (K) Heatmap of indicated genes reported to be induced upon bacteria binding to IECs are increased in *Ire1α^-/-^Ireβ^-/-^* IECs. Transcript per million (TPM) values for each gene are row normalized to z-scores. Each column represents a biological replicate belonging to the indicated genotypic group.

We then isolated IECs for RNA sequencing (RNA-Seq) from 6 biological replicates for each of *WT*, *Ire1α^-/-^, and Ire1β^-/-^* and 5 biological replicates for *Ire1α^-/-^Ire1β^-/-^* double deleted mice. After processing and QC of the data, expression of 15,641 genes was detected in these samples and used for downstream analysis (Fig. S3A-D). Principal component analysis (PCA) demonstrated clustering of biological replicates within each genotypic group and separation of *Ire1α^-/-^Ire1β^-/-^* mice from *WT,* intestine-specific *Ire1α^-/-^* and *Ire1β^-/-^* mice along PC1 that accounted for 27% of variance in the data (Fig. S3E). Importantly, no significant changes in gene expression were detected in IECs from intestine-specific *Ire1α^-/-^* or *Ire1β^-/-^* mice compared to the *WT* group, consistent with histological observations (Fig. S3F, G). In contrast, we observed extensive mRNA changes in IECs from double deleted *Ire1α Ire1β* mice, where 225 genes were up-regulated and 477 were down-regulated (fold-change ≥ 2.0, BH-adjusted p-value < 0.05) in *Ire1α^-/-^Ire1β^-/-^* versus *WT* mice (Fig. S3H). Expression of mRNAs encoding defensins and LYZ were decreased in double deleted *Ire1α^-/-^Ire1β^-/-^* IECs but not in single deleted *Ire1α^-/-^* or *Ire1β^-/-^* IECs (Fig. 3G). Matrix metalloproteinase 7 (MMP7) and Angiogenin 4 (Ang4) were also significantly decreased in double deleted *Ire1α^-/-^Ire1β^-/-^* IECs (Fig. 3G). MMP7 is responsible for activating defensins, while Angiogenin 4 (Ang4) is an antimicrobial constituent secreted from Paneth cells. Additionally, complement expression was decreased only in double deleted *Ire1α^-/-^Ire1β^-/-^* IECs (Fig. 3H). Consistent with this finding, mRNAs encoding innate immune responses were also uniquely reduced in double deleted *Ire1α^-/-^Ire1β^-/-^* IECs (Fig. 3I). In contrast, mRNAs encoding functions in enteritis/colitis were uniquely increased in double deleted *Ire1α^-/-^Ire1β^-/-^* IECs (Fig. 3J). mRNAs induced upon microbiome binding to IECs were significantly upregulated only in double deleted *Ire1α^-/-^Ire1β^-/-^* IECs (Fig. 3K). Taken together, mRNAs encoding components of intestinal immune defense were extensively decreased, while mRNAs that exert pathogenic roles were significantly increased in double deleted *Ire1α^-/-^Ire1β^-/-^* IECs.

### Deletion of *Ire1α* and *Ire1β* disrupts the colonic epithelium

We then evaluated the impact on the colon of double deletion of *Ire1α* and *Ire1β*. Architectural distortion was observed in both young and aged adult double deleted *Ire1α^-/-^Ire1β^-/-^* mice (Fig. 4A; Fig. S4A). Alcian blue positive cells were decreased in double deleted *Ire1α^-/-^Ire1β^-/-^* colons but were not altered in single deleted mice (Fig. 4B; Fig. S4B). Consistent with this observation, mucin secretory granules were infrequently observed by transmission electron microscopy (TEM) in double deleted *Ire1α^-/-^Ire1β^-/-^* colons (Fig. 4C). These changes were not found in colons from mice with single deletion of either *Ire1α* or *Ire1β* (Fig. S4A, B&C). We further identified increased numbers of Ki67 positive cells in double deleted *Ire1α^-/-^Ire1β^-/-^* mice (Fig. 4D), suggesting increased proliferation. In addition, spontaneous polypoid lesions were grossly visible in colons from 4 of 32 double deleted *Ire1α^-/-^Ire1β^-/-^* aged mice but not in *WT* and single deleted *Ire1α^-/-^* or *Ire1β^-/-^* mice (Fig. 4E & Fig. S4D). We also observed rectal prolapse in 10 of 32 *Ire1α^-/-^Ire1β^-/-^* mice (Fig. S4E), and histologic examination demonstrated severe inflammation and residual intestinal glands (Fig. S4E). These observations demonstrate severe colonic epithelial disruption upon double deletion of *Ire1α* and *Ire1β*.

**Fig. 4.**
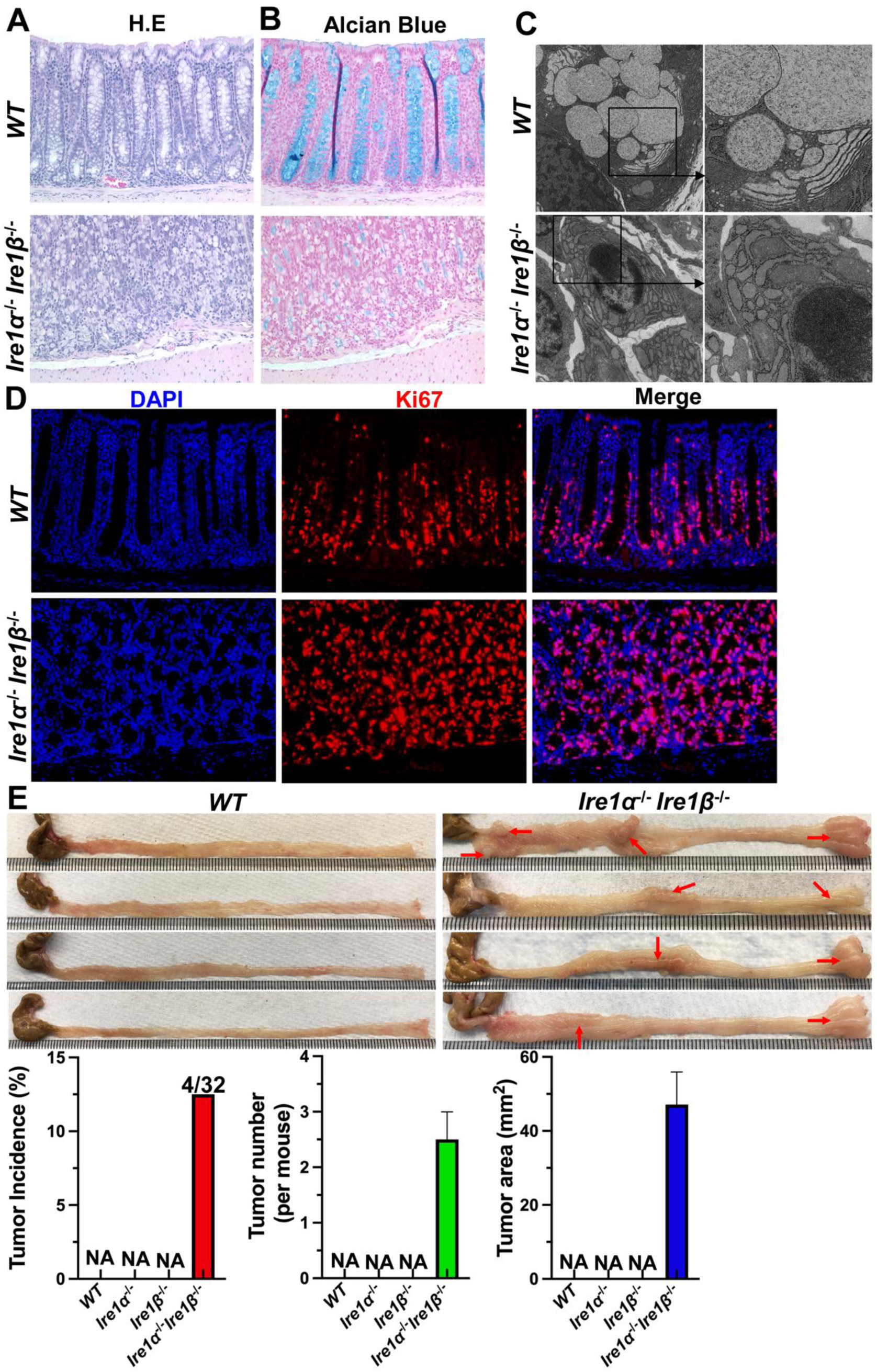
*Ire1α^-/-^Ireβ^-/-^* mice display colonic mucosal abnormality. (A) H&E staining shows colon architecture distortion in aged *Ire1α^-/-^Ireβ^-/-^* mice. (B) Alcian blue staining shows few positive cells in aged *Ire1α^-/-^Ireβ^-/-^* mice. (C) TEM shows absence of secretory particles in *Ire1α^-/-^Ireβ^-/-^* mice but not in age matched WT mice. (D) Immunofluorescence shows increased Ki67 positive cells in *Ire1α^-/-^Ireβ^-/-^* mice compared to age matched WT mice. (E) Spontaneous colorectal tumors occur in *Ire1α^-/-^Ireβ^-/-^* mice but not in age matched WT, *Ire1α^-/-^* or *Ireβ^-/-^* mice.

### Deletion of either *Ire1α* or *Ire1β* increases colonic tumorigenesis

The absence of a spontaneous intestinal phenotype with IEC-specific *Ire1α* deletion led us to ask whether a phenotype upon *Ire1α* deletion would emerge following AOM-DSS challenge. First, we challenged WT *(Ire1α^+/+^)* and intestine-specific *Ire1α^-/-^* mice with one AOM injection (12.5mg/Kg) at 6-8wks of age followed by two cycles of 7d 2.5% DSS treatment with an interval of 14d. Tumor formation was increased in AOM-DSS treated *Ire1α^-/-^* mice compared to *WT (Ire1α^+/+^)* mice (P<0.01, Fig. 5A). To extend this observation, we crossed *APC*^Min^ C57BL/6J mice with *IRE1α^fl/fl^* Villin Cre C57BL/6J mice to generate *APC*^Min^*Ire1α^fl/fl^* Villin Cre (*APC*^Min^*Ire1α^-/-^*) and *APC*^Min^*Ire1α^fl/fl^* (*APC*^Min^*Ire1α^+/+^*) mice (20). We found increased colonic tumors in *APC*^Min^ *Ire1α^-/-^* mice compared to *APC*^Min^ *Ire1α^+/+^* mice (P<0.05, Fig. 5B). Collectively, these data indicate that IEC-specific *Ire1α* deletion increases colonic tumorigenesis.

**Fig. 5.**
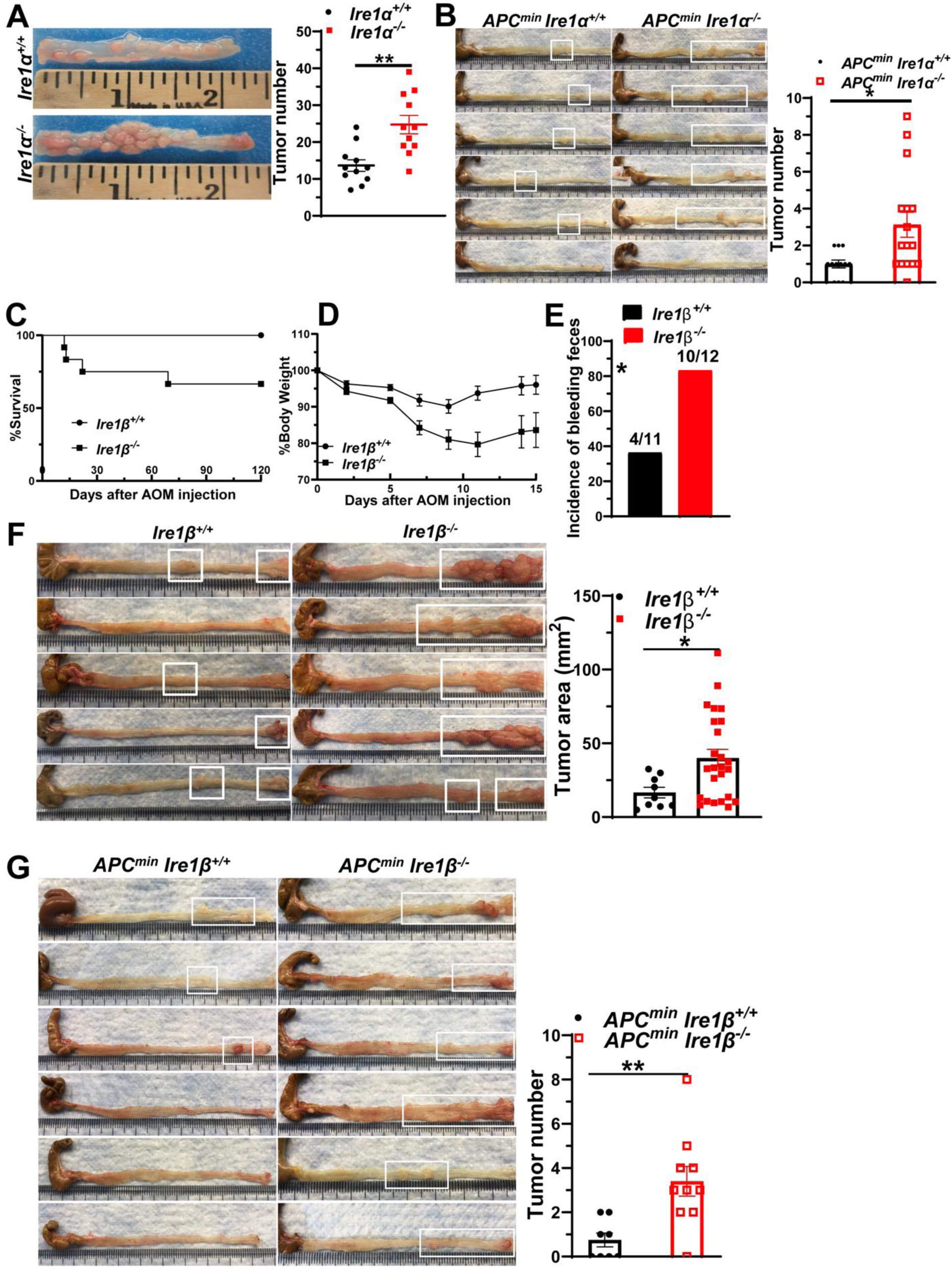
*Ire1α* or *Ire1β* deficiency increases CRC in multiple murine models. (A) *Ire1α^+/+^*(WT) and *Ire1α^-/-^* mice were challenged with one AOM injection (12.5mg/Kg) at 6-8wks of age followed by two 7d cycles of 2.5% DSS treatment with an interval of 14d. Colorectal tumor numbers were measured at 4 mon after AOM injection and are shown as mean ± SE (n=11 mice). ** P=0.0012. (B) *APC^min^ Ire1α^+/+^* and *APC^min^ Ire1α^-/-^* mice were harvested and colorectal tumors were counted at ∼16 wks of age. * P=0.0147 (n= 12 mice for *APC^min^ Ire1α^+/+^* & 16 mice for and *APC^min^ Ire1α^-/-^*). *Ire1β^+/+^* (WT) and *Ire1β^-/-^* mice were treated with AOM (12.5mg/kg) at 6-8 wks of age followed by 7 days of 2% DSS treatment. (C) Survival curve P=0.0398. (D) Body weight changes. (E) Incidence of bleeding feces upon DSS treatment, P=0.0361, *Ire1β^+/+^* n=11 mice, *Ire1β^-/-^* n=12 mice. (F) Colorectal tumors measured 4 mon after AOM injection. *Ire1β^+/+^* n=11 mice, *Ire1β^-/-^* n=8 mice, each dot in the graph represents one tumor mass. Data are shown as means ± SE *P=0.025. (G) *APC^min^ Ire1β^+/+^* and *APC^min^ Ire1β^-/-^* mice were harvested and colorectal tumors were counted at ∼16 wks of age. *APC^min^ Ire1β^+/+^* n=8 mice, and *APC^min^ Ire1β^-/-^* n=10 mice. Each dot in the graph represents one mouse **P=0.0045.

To examine the role of *Ire1β* in a similar scenario, we injected *WT* (*Ire1β^+/+^*) and *Ire1β^-/-^* mice with AOM (12.5 mg/kg), followed by one cycle 2% DSS in drinking water. We observed decreased survival (Fig. 5C) and severe body weight loss in *Ire1β^-/-^* mice with a nadir of 79% ± 6.1 (mean ± SE) compared to *WT* (*Ire1β^+/+^*) mice with a nadir of 91%± 3.8 (mean ± SE) (Fig. 5D). Additionally, 83.3% (10/12) of *Ire1β^-/-^* mice exhibited frank fecal blood but only 36.6% (4/11) of *WT* (*Ire1β^+/+^*) mice exhibited fecal blood (Fig. 5E). These findings indicate that DSS exposure produces more severe colitis in *Ire1β^-/-^* mice, consistent with previous findings (10). We then measured colon tumor formation in *Ire1β^-/-^* mice. The tumor area was increased in the colons of *Ire1β^-/-^* mice compared to WT mice (p< 0.05, Fig. 5F). These data indicate that the *Ire1β^-/-^* colon is more prone to tumor formation in the setting of AOM-DSS induced colitis associated CRC, consistent with more severe colitis in *Ire1β^-/-^* mice. We also generated *APC^min^Ire1β^-/-^* mice and observed more colon tumors compared to *APC^min^Ire1β^+/+^* mice (p<0.01 Fig. 5G). Collectively, these data show *Ire1β* deletion increases colonic tumorigenesis, and, combined with the findings above, suggest that intestinal *Ire1α* and *Ire1β* individually function as CRC tumor suppressor genes.

### *Ire1α* or *Ire1β* deletion increases colon stem cell survival and growth

To test whether *Ire1α^-/-^* ISCs exhibit a clonal survival advantage, we backcrossed *Lgr5-eGFP-Cre^ER^-Lox-STOP-Lox-tdtomato* mice with *Ire1α^fl/fl^* mice to generate Lgr5-eGFP-Cre^ER^-Lox-STOP-Lox-tdtomato-*Ire1α^fl/fl^* (*eGFP-tdtomato-Ire1α^-/-^*) and Lgr5-EGFP-Cre^ER^-Lox-STOP-Lox-tdtomato-*Ire1α*^+/+^ (*eGFP-tdtomato-Ire1α*^+/+^) mice. All Lgr5 positive cells expressed eGFP and tdtomato expression that was induced at 20d after low dose Tamoxifen (Tam) (0.15mg/mouse) injection (Fig. 6A). In this model, all non-induced ISCs located at the base of the crypt were labeled as green (Fig. 6A, left image), whereas newly generated Lgr5^+^ cells were labeled yellow, a signal from the emerging expression of tdtomato and eGFP (Fig. 6A, right image). Using this model, we compared the *eGFP-tdtomato-Ire1α^-/-^* ISC average clone size with their *eGFP-tdtomato-Ire1α^+/+^* littermates. Mice were injected with low dose Tam (0.15mg) combined with 7d of 2.5% DSS challenge. The average clone size with 7d DSS (2.5%) challenge was unchanged in small intestinal clones but increased in colonic crypts from *Ire1α^-/-^* mice (Fig. 6B, P< 0.05). These data support the notion that *Ire1α* deletion confers a selective survival advantage to colonic ISCs.

**Fig. 6.**
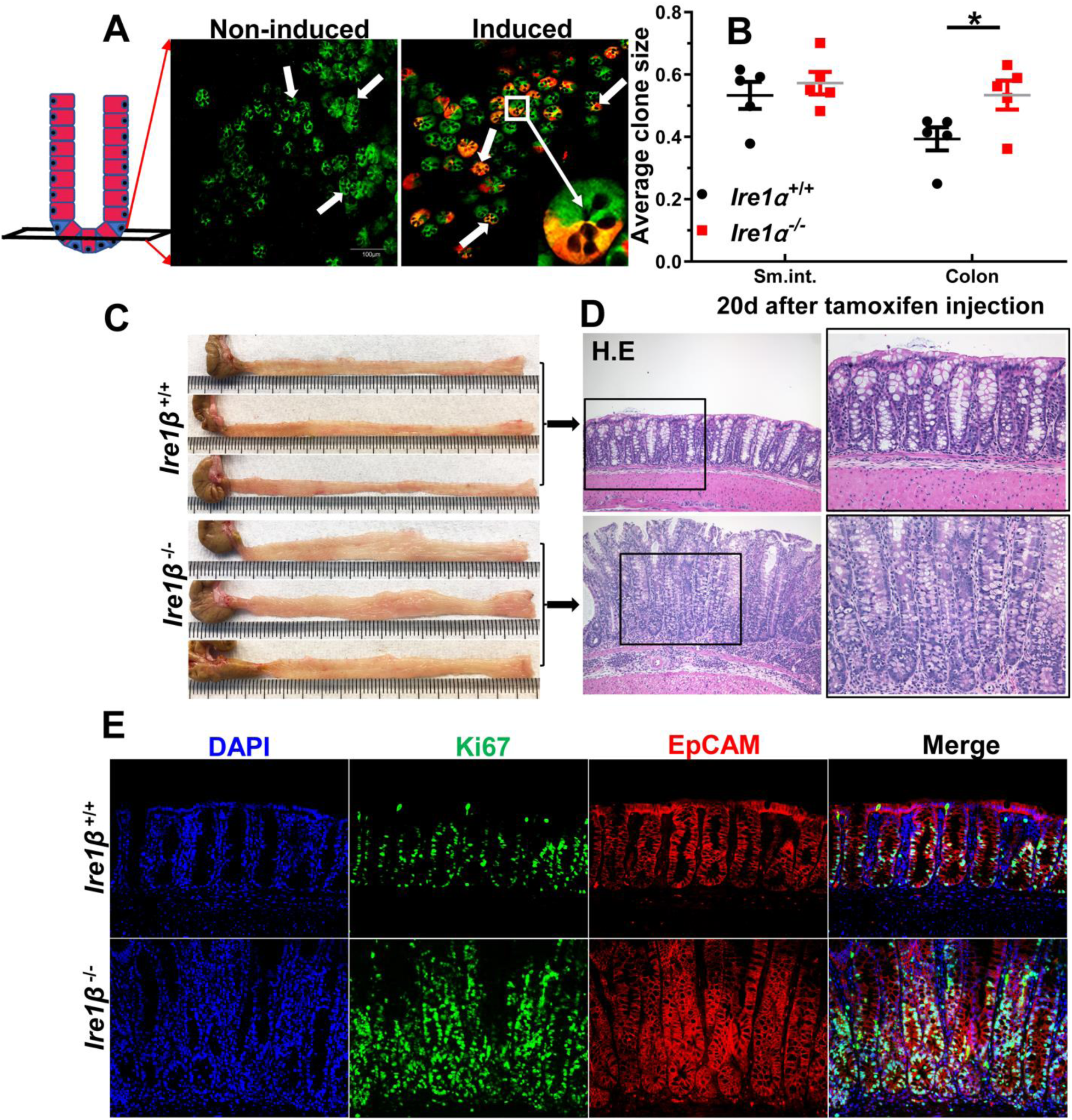
*Ire1α* or *Ire1β* deletion increase intestinal stem cell survival. (A) Confocal images show small intestinal crypts of eGFP-tdtomato-*Ire1α^+/+^* mice without or with 0.15mg Tam induction (20d after injection). (B) Average clonal sizes of eGFP-tdtomato-*Ire1α^+/+^* and eGFP-tdtomato-*Ire1α^-/-^* ISCs, calculated at 20d after Tam injection with 2.5% DSS challenge are shown as means ± SE (n=5). *P=0.045 (C) Images show morphology of *Ire1β^+/+^* and *Ire1β^-/-^* colons harvested 21 days after AOM injection and 15 days after 2% DSS treatment. (D) H&E staining of *Ire1β^+/+^* and *Ire1β^-/-^* colons is shown: 100x magnification for left two panels and 200x magnification for right two panels. (E) Immunofluorescence micrograph of Ki67 and Epcam on *Ire1β ^+/+^* and *Ire1β ^-/-^* colons are shown (n=6 mice/group). 200x magnification.

To test if ISC regeneration is also enhanced upon *Ire1β* deletion, we treated *Ire1β^-/-^* mice with 12.5mg/kg AOM followed by 6d 2% DSS. After 15d of drinking water washout, colons were harvested for immunofluorescence to visualize Ki67 and Ep-CAM double positive cells. We observed macroscopic disruption of the colonic mucosa in *Ire1β^-/-^* mice as early as 3 weeks after AOM injection (Fig. 6C), along with epithelial hyperplasia and hypertrophy as shown by H&E staining (Fig. 6D). Correspondingly, we detected enhanced Ki67 and Epcam double positive proliferating epithelial cells from colons of *Ire1β^-/-^* mice (Fig. 6E). These data together suggest there is increased colonic ISC regeneration upon *Ire1β* deletion.

### *Ire1α* and *Ire1β* regulate different sets of genes to maintain intestinal homeostasis

As noted above, we observed enhanced tumorigenesis in AOM/DSS treated *Ire1α^-/-^* mice, yet disease severity in *Ire1α^-/-^Ire1β^-/-^* mice was both more severe and more extensive than that reported in IEC-specific *Xbp1* deleted mice(12). Those observations together suggest that *Ire1β* may regulate distinct downstream pathways from *Ire1α*. To test this notion, we analyzed the expression of a set of mRNAs downstream of XBP1s using the RNA-Seq data. We found that expression of mRNAs was distinct between *Ire1α^-/-^* single deleted and *Ire1β^-/-^* single deleted IECs. Significantly, those mRNAs that were significantly decreased in *Ire1α^-/-^* IECs were not reduced in *Ire1β^-/-^* IECs (Fig. 7A). This provides evidence that gene regulation through IRE1α and IRE1β is different, and that *IRE1β* does not obviously splice XBP1 mRNA.

**Fig. 7.**
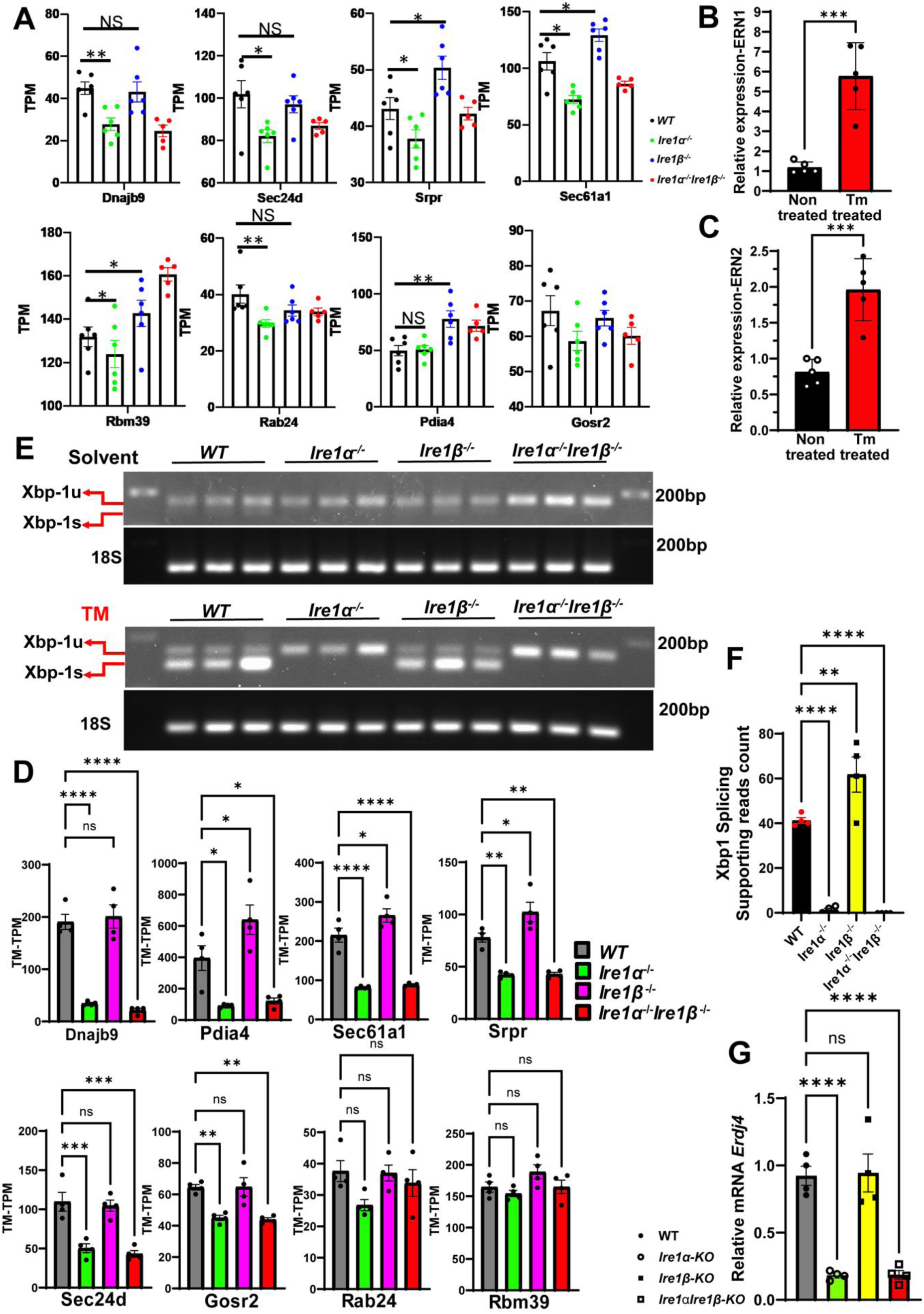
*IRE1β* does not detectably splice *Xbp1* mRNA. (A) Expression of indicated *Xbp1s* target genes summarized from RNA-Seq data from indicated IECs without Tm treatment. (B&C) Five 12-weeks-old WT mice were injected with 1mg/kg of Tunicamycin or solvent, IECs were harvested 12 hours later and RNA was extracted. Ern1 (*Ire1α*) and *Ern2* (*Ire1β*) were measured by RT-PCR. (D) Expression of indicated *Xbp1s* target genes summarized from Tm-treated IECs RNA-Seq data. (E) 10-14 weeks old mice were injected with 1mg/kg of Tunicamycin or solvent and IECs were harvested 12 hours later. RNA was extracted and *Xbp1* splicing was measured by RT-PCR and visualized by 2% Agarose gels (N=3 mice for each group). (F) The Read-split-Run, an improved bioinformatic pipeline, was used to analyze Tm-treated IECs RNA-Seq data to detect *Xbp1* mRNA splicing. The supporting read counts of the detection of non-canonical spliced regions of *Xbp1* mRNA is summarized (n=4 mice for each group). (G) Expression of Erdj4 from Tm-treated indicated mice was evaluated by RT-qPCR. (n=4 mice for each group).

*Ire1α* and *Ire1β* mRNAs were significantly increased in IECs from WT mice following induction of ER stress using a 12-hour, non-lethal dose (1mg/kg) of tunicamycin treatment, (Fig. 7B&C). We then treated *WT*, *Ire1α^-/-^, Ire1β^-/-^* and double deleted *Ire1α^-/-^Ire1β^-/-^* mice with the same dose of tunicamycin, to test whether IRE1β mediates *Xbp1u* splicing following induction of ER stress. IECs were collected 12 hours after treatment, and RNA was extracted for subsequent RNA sequencing and analyses (Fig. S5A-E). We examined the expression of gene sets downstream of XBP1s using RNA-Seq data for IECs from mice treated with Tm, following the same approach used for RNA-Seq data for IECs from untreated mice. Consistently, we found that expression of these mRNAs differed between *Ire1α^-/-^* and *Ire1β^-/-^* IECs and mRNAs that exhibited decreased expression in *Ire1α^-/-^* IECs, which are known targets of XBP1s, did not show a similar decrease in *Ire1β^-/-^* IECs (Fig. 7D). These data suggest that IRE1β does not detectably splice XBP1 mRNA, even following tunicamycin treatment and induction of ER stress.

We then analyzed IEC mRNA from new cohorts of mice following tunicamycin treatment and conducted gel analysis of *Xbp1s* to further confirm that IRE1β does not detectably splice *Xbp1u* mRNA. As shown in Fig 7E, *Xbp1s* mRNA was barely detectable in IECs without tunicamycin treatment. Only a faint *Xbp1* RT-PCR product was visible in WT and *Ire1β^-/-^* IECs and no band was detected in *Ire1α^-/-^* IECs, where IRE1β mRNA expression was preserved (Fig. 7E). In WT and *Ire1β^-/-^* IECs, *Xbp1s* mRNA bands were readily detected upon tunicamycin stimulation. However, *Xbp1s* was not detected in *Ire1α^-/-^* IECs, even under conditions (ie tunicamycin treatment) *Xbp1* splicing was readily observed in WT and *Ire1β^-/-^* IECs (Fig. 7E).

We then asked whether there is a novel non-canonical splicing target gene unique to *Ire1β,* using an improved algorithm, Read-Split-Run, to identify potential non-canonical splice junctions (21)., using RNA-Seq data from tunicamycin-treated *WT*, *Ire1α^-/-^, and Ire1β^-/-^* and double deleted *Ire1α^-/-^Ire1β^-/-^* IECs. However, no distinctive non-canonical splicing junctions specific for IRE1β were identified. We detected non-canonical splice junctions of *Xbp1s* in both *WT* and *Ire1β^-/-^* IECs but not in IECs from *Ire1α^-/-^* and double deleted *Ire1α^-/-^Ire1β^-/-^* mice (Fig. 7F). Those findings align with the expression of *Erdj4*, a downstream target gene for *XBP1s*, which was decreased in IECs from *Ire1α^-/-^ and* double deleted *Ire1α^-/-^Ire1β^-/-^* mice as analyzed by RT-qPCR (Fig. 7G). Taken together, these results reinforce the notion that IRE1β does not splice *Xbp1u*, and that suggest that suppression of intestinal tumorigenesis via IRE1α and IRE1β occur through different mechanisms.

Next, we conducted GSEA analysis on the RNA-Seq data by mapping differential mRNA profiles between tunicamycin-treated *Ire1α^-/-^* vs WT, as well as between *Ire1β^-/-^* vs WT, to the 187 gene sets listed in the GSEA C6 oncogenic signature category. We found that 23 out of these 187 gene sets were upregulated in *Ire1β^-/-^* IECs (with a p value<0.05 and a False Discovery Rate<10%). In contrast, none of these 23 gene sets were observed to be upregulated in *Ire1α^-/-^* IECs, underscoring the distinct mechanisms utilized by *Ire1α* and *Ire1β*, with_distinct oncogenic pathways enhanced exclusively in *Ire1β^-/-^* IECs (Fig. S5F). We observed a gene set downstream of mTORC1 to be increased upon *Ire1β* deletion, suggesting that suppression of mTORC1, a growth promoting signaling pathway, may be utilized by IRE*1β* to maintain intestinal homeostasis and suppress CRC (Fig. 8A). The upregulation of this gene set in *Ire1β^-/-^* IECs was further confirmed through RT-qPCR analysis of mRNAs from tunicamycin-treated WT, *Ire1α^-/-^*, and *Ire1β^-/-^* IECs (Fig. 8B). Because tunicamycin stimulation was not used to induce CRC, we treated WT and *Ire1β^-/-^* mice with 2% DSS and WT and *Ire1α^-/-^* mice with 2.5% DSS for one week to ask whether a corresponding pattern of mRNAs might be induced following inflammatory colonic injury. Those experiments revealed increased mRNA expression in the mTORC1 gene set in DSS-treated *Ire1β^-/-^* IECs but not in *Ire1α^-/-^* IECs (Fig. 8C&D). These data reinforce the suggestion that the augmentation of CRC in *Ire1β^-/-^* mice may be associated with an increase in gene expression downstream of mTORC1.

**Fig. 8.**
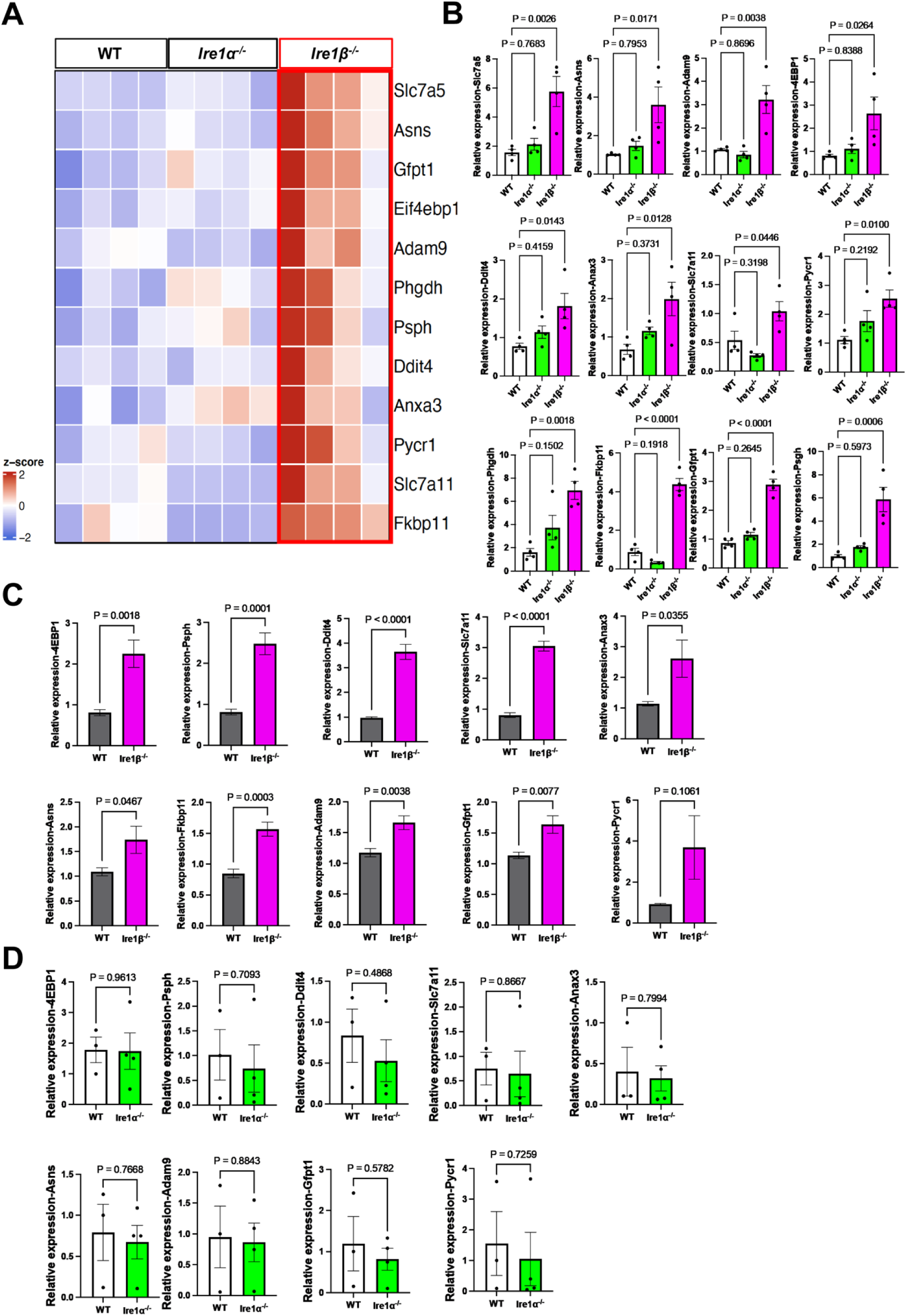
Gene set enrichment analysis downstream of mTORC1 that are increased in *Ire1β*^-/-^ IECs. (A) Heatmap of selected genes from Tm-treated IECs RNA-Seq data shows expansion of a genes downstream of mTORC1 that are increased in *Ire1β*^-/-^ IECs. (n=4 mice for each group) (B) RT-qPCR evaluation is shown for the gene set downstream of mTORC1 in Tm-treated IEC RNA (n=4 mice for each group). (C&D) RT-qPCR analysis of the gene set downstream of mTORC1 in DSS-treated colon IECs mRNA is shown (n=5 mice for each group in C and n=4 mice for *Ire1α^-/-^* and n=3 mice for WT group in D).

### Increased *IRE1α* and *IRE1β* expression correlates with improved survival in CRC patients

We then asked whether our observations of enhanced CRC upon *Ire1α* or *Ire1β* deletion in mouse models might be recapitulated in patients with CRC. We analyzed *IRE 1β* mRNA expression from seven Oncomine datasets, a total 447 CRC patients and 192 normal control samples. We found that *IRE1β* expression was decreased in human CRC tissue compared to normal tissue (P<0.001, Fig. 9A), demonstrating an association between decreased *IRE1β* and the occurrence of CRC. However, *IRE1β* expression in CRC patients was not uniform. This uneven distribution led us to study the correlation of patient survival with *IRE1β* expression. From three PronoScan datasets, from a total of 126 cases with higher *IRE1β* expression and 300 cases with lower levels, we found that patients with higher *IRE1β* mRNA expression exhibited improved survival compared to patients with lower levels (Fig. 9B, P<0.05). Although we did not observe higher *IRE1α* expression in CRC when compared to normal controls, we did find that CRC patients with higher *IRE1α* expression survive longer than those with lower levels (Fig. 9C). These findings suggest concordance with our observations in mice and with patient outcomes in CRC, implying that stratifying CRC patients by *IRE1α* or *IRE1β* expression might represent a potential prognostic biomarker.

**Fig. 9.**
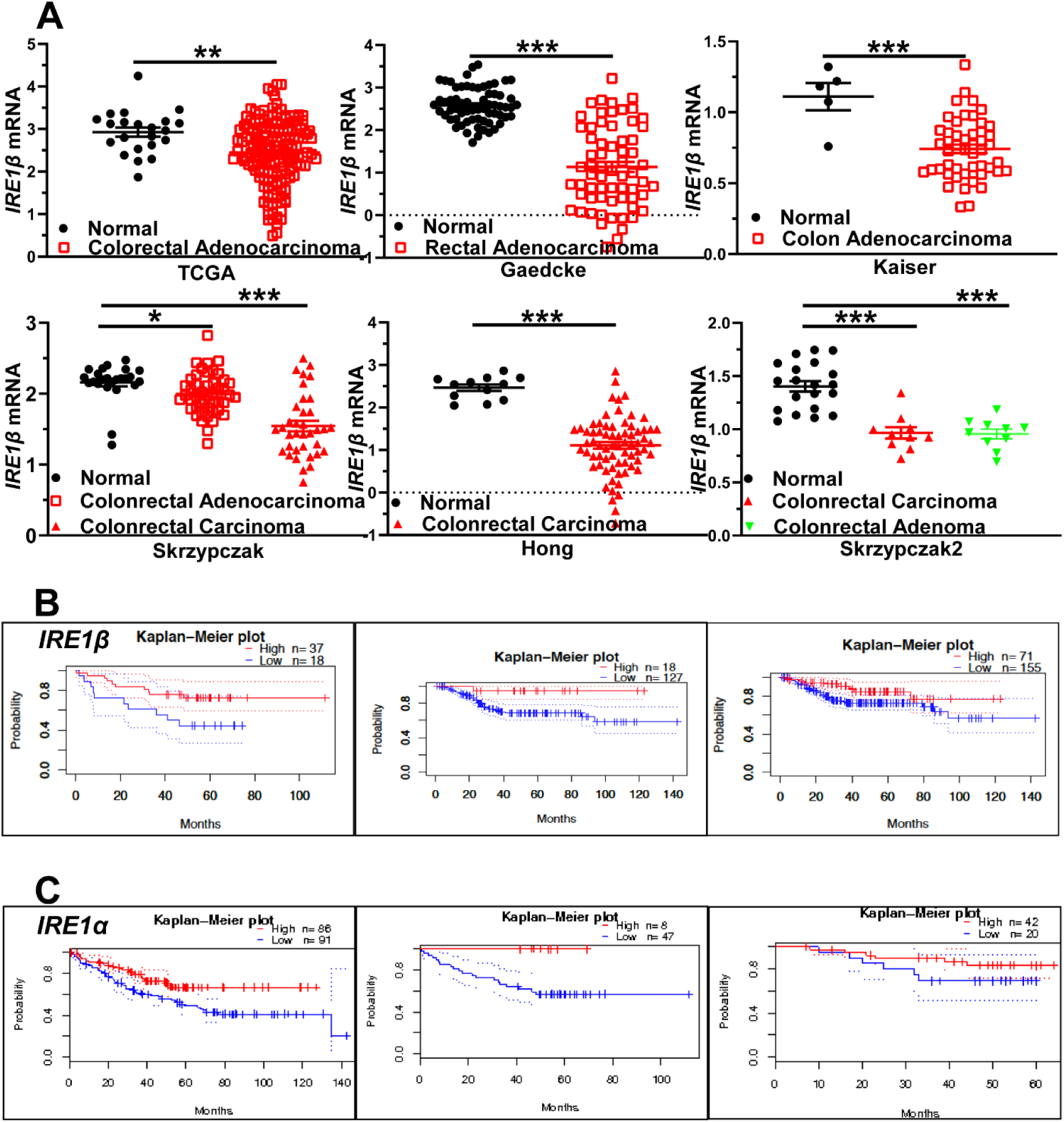
Decreased expression of *IRE1β* in CRC patients and increased *IRE1α* and *IRE1β* expression predicts better survival in CRC patients. (A) *IRE1β* mRNA from colorectal adenocarcinoma, carcinoma, adenoma patients and healthy control tissue is shown. All data were obtained from the oncomine database. Panel 1, TCGA dataset, n= 22 VS162 (healthy VS adenocarcinoma) P=0.0045; Panel 2, Gaedcke dataset, n= 65 VS 65(healthy VS adenocarcinoma) p<0.0001; Panel 3, Kaiser dataset n= 5 vs 49 and (healthy VS adenocarcinoma) p=0.0005; Panel 4, Skrzypczak dataset n=24 vs 45 (healthy VS adenocarcinoma) p=0.027, n=24 vs 36 (healthy VS carcinoma) p<0.0001; Panel 5, Hong dataset n= 12 vs 70 and (healthy VS carcinoma) p<0.0001; Panel 6, Skrzypczak2 dataset n= 20 vs 10 (healthy VS carcinoma) p<0.0001, n= 20 vs 10 (healthy VS adenoma) p<0.0001; Data are shown as means ± SE. (B) Kaplan-Meier plots show survival for CRC patients with higher (red) or lower (blue) *IRE1β* levels. Panel 1, VMC dataset, n_high_=37, n_low_=18, P=0.0366. Panel 2 MCC dataset, n_high_=18, n_low_=127, P=0.030797; Panel 3 Melbourne dataset, n_high_=71 n_low_=155, P=0.049552. (C) Kaplan-Meier plots of survival for CRC patients with higher (red) or lower (blue) *IRE1α* levels. Panel 1 (MCC dataset, n_high_=86, n_low_=91, P=0.010), Panel 2 (VMC dataset, n_high_=8, n_low_=47, P=0.038), Panel 3 (Berlin dataset, n_high_=42, n_low_=20, P=0.169). Datasets were obtained from the prognoscan database.

## Discussion

IRE1 is the most conserved UPR signaling arm that exists in all cells of eukaryotes from yeast to mammals. However, in addition to Ire1α, epithelial cells of the gut and the lung have evolved a second paralogue, Ire1β. IRE1α RNase specifically cleaves the double hairpin motif of unspliced XBP1 mRNA, leading, after ligation of the exons by the RTCB tRNA ligase (7), to splice XBP1 (XBP1s) mRNA. Other work showed IRE1α RNase activity promotes regulated IRE1-dependent decay (RIDD) (22). In contrast, a cleavage target for IRE1β has remained elusive despite observations that it cleaves MTTP mRNA and 28S RNA (19, 23) and exhibits weak XBP1 splicing activity (24). Here, we show that IRE1α and IRE1β both maintain intestinal homeostasis and suppresses tumorigenesis, although each operates through a different mechanism (Fig. 10).

**Fig. 10.**
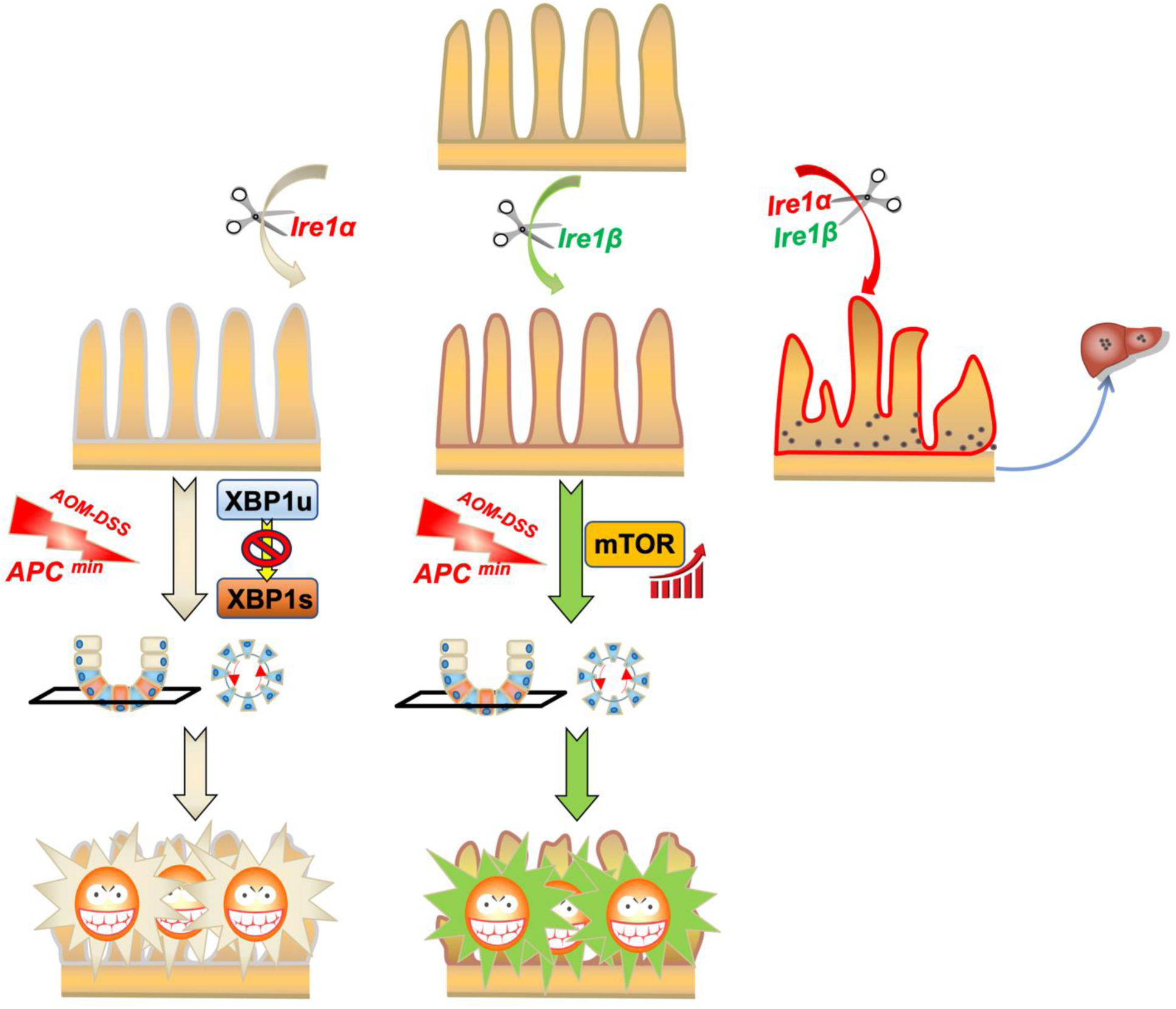
Graphic abstract depicts the importance of IRE1α and IRE1β in maintaining intestinal homeostasis and suppressing intestinal tumor formation.

Intestinal homeostasis remained well balanced, with no apparent damage when either *Ire1α* or *Ire1β* was deleted from IECs, without either an external (*i.e.,* DSS) or further genetic (double deletion) insult. Our current findings regarding the basal phenotype in intestine-specific *Ire1α* null mice failed to replicate other reports where deletion of *Ire1α* induced spontaneous colitis and progressive mortality (15). Differences in the microbial communities or other environmental factors may have contributed to the discrepant baseline phenotypes observed in our line of intestine-specific *Ire1α* null mice compared to the line generated by Zhang and colleagues (15) and will require further study to resolve. In contrast, epithelial homeostasis was disrupted and extensive injury observed in both small intestine and colon only when both IEC *Ire1α* and *Ire1β* were deleted. However, in both the colitis (AOM-DSS)-induced and the *APC^min^* genetic insult models, homeostasis was disrupted and CRC was enhanced when either *Ire1α* or *Ire1β* was lost. The augmentation of CRC in intestine-specific *Ire1α*^-/-^ single deleted mice is likely attributable to the absence of Xbp1s, because IEC loss of Xbp1s was observed to augment intestinal tumorigenesis (13). As alluded to above, we found that IRE1β does not detectably splice *Xbp1u*-mRNA, suggesting the observed increased tumorigenesis upon *Ire1β* deletion may reflect increased expression mTORC1-dependent genes.

We evaluated the expression patterns of *Ire1α* and *Ire1β* in IECs and found that they are both highly expressed in goblet cells, Paneth cells, transit amplifying cells, and ISCs. The co-expression of these two molecules in diverse cell types suggests that under some (but perhaps not all (17) circumstances), the absence of either IRE1 paralogue can be compensated by the remaining paralogue, suggesting they provide complementary but distinct functions to preserve intestinal homeostasis. We did not examine the role of microbial signaling in our double deleted *Ire1α Ire1β* IECs, and it is conceivable that under germfree conditions a different injury phenotype may emerge, as observed in mice with *Ire1β* deletion (17). However, our findings suggest that while no overt injury was observed in chow fed mice with intestine-specific *Ire1α* mice or in germline *Ire1β^-/-^* mice at baseline, we found enhanced tumorigenesis following carcinogen-DSS stimulation or the APC^min^ genetic challenge. Those findings suggest that a single IRE paralogue is insufficient in mitigating the tumorigenic phenotypes observed.

The simultaneous expression of *Ire1α* and *Ire1β* in subsets of IECs (24) may explain the spontaneous damage observed in the intestinal tract when these paralogs are both depleted. It is noteworthy that goblet cells and Paneth cells co-express both *Ire1α* and *Ire1β*, since these cells secrete various mucins, defensins, and LYZ, which are essential to protect the intestinal tract (1, 24). Further studies are needed to clarify how these two molecules cooperate to maintain protein folding and secretion in goblet and Paneth cells and the cell-specific impact of protein misfolding in the ER (25, 26). As noted above, previous studies demonstrated IRE1β−dependent splicing of *Xbp1*-mRNA in an *in vitro* culture system (17, 24). In contrast, we did not detect any change in *Xbp1u* mRNA splicing in *Ire1β^-/-^* IECs *in vivo* (Fig 7). Our studies reflect the in-vivo situation, however, rather than colonoids but we used the same line of *Ire1β^-/-^* mice (10). In that earlier study, the IRE1 inhibitor 4μ8C was applied to an *in vitro* colon crypt culture system containing both WT and *Ire1β*^-/-^ crypts (17). Notably, *Xbp1s* mRNA expression was found to be higher in *Ire1β*^-/-^ crypts treated with 4μ8C compared to *Ire1β*^-/-^ crypts without treatment, where IRE1α was still present (17). Previously, 4μ8C was described as an IRE1 inhibitor (27), and this inhibitor is commonly employed to target the IRE1α-XBP1s pathway (28). However, in the absence of evidence that 4μ8C effectively inhibits IRE1β, we suggest that 4μ8C is not an ideal choice for assessing whether IRE1β splices *Xbp1* mRNA. For example, the expression of spliced *Xbp1* mRNA was observed to increase upon 4μ8C treatment in *Ire1β*^-/-^ crypts (27), a situation where IRE1α-driven splicing was suppressed (29). We also observed that mice with *Ire1α* and *Ire1β* double deletion display extensive disruption of the intestinal tract that is more profound than the damage observed in *Xbp1* deleted mice(12), which further suggests that IRE1β performs additional functions, besides splicing *Xbp1* mRNA, to maintain intestinal homeostasis. These findings should also be interpreted with the background knowledge that IRE1a and IRE1b function as heterooligomers in addition to their individual role in mucosal homeostasis (30).

Consistent with our findings in mice, we found that *IRE1β* expression decreased in multiple datasets from patients with CRC and we further observed that higher levels of *IRE1α* or *IRE1β* mRNA expression correlated with improved survival in patients with CRC. These observations indicate that analysis of *IRE1α* and *IRE1β* expression in patients with CRC might assist in stratifying patients for therapy and for predicting prognosis. It remains unknown whether there are mutations in *IRE1α* or *IRE1β* in patients with CRC, but large-scale screening of patients to identify mutations in these two genes would be potentially valuable in a clinical setting. Expression of *IRE1α* or *IRE1β* may not only predict the prognosis of CRC patients but also may provide insight regarding the sensitivity of mTOR inhibitor treatment (31). Rapalogs, mTOR inhibitors such as Everolimus (Afinitor) and Temsirolimus (Torisel), are FDA-approved medicines used for the treatment of various cancers (32). Our data support the possibility that patients with either reduced levels of *IRE1β* expression or that harbor inactivating *IRE1β* mutations may be more sensitive to Rapalog treatment. This notion is presently being tested in mice. In addition, it was reported that the FDA-approved medicines including methotrexate, fludarabine phosphate and sunitinib can inhibit IRE1 activity (33–35). Those findings, together with our results, raise concerns about whether these treatments might influence the development of progression of CRC.

## Materials and Methods

### Mice

*Ire1α* ^fl/fl^ C57BL/6J, were originally generated by our lab (36). We crossed this line with *Villin* Cre to generate intestine epithelial cell specific deleted *Ire1α*^fl/fl^ *Villin* Cre (*Ire1α^-/-^*) and *Ire1α* ^fl/fl^ (*Ire1α^+/+^ or* WT) littermate control. We crossed *APC^min^* C57BL/6 (JAX, # 002020) with *Ire1α*^fl/fl^ *Villin* Cre to get *APC^min^ Ire1α*^fl/fl^ *Villin* Cre (*APC*^Min^ *Ire1α^-/-^*) and littermate control *APC* ^min^ *Ire1α*^fl/fl^ (*APC*^Min^ *Ire1α^+/+^*). Germline *Ireβ^-/-^* mice were kindly provided as a gift from Dr. David Ron in which exons 12 and 13 that encode the transmembrane domain were deleted. We crossed this line with *APC^min^* C57BL/6 (JAX, #002020) to obtain *APC^min^ Ire1β^+/-^* and l*Ire1β^+/-^* mice. We mated *APC^min^ Ire1β*^+/-^ mice with *Ire1β^+/-^* mice to obtain *APC ^min^ Ire1β^-/-^* and littermate *APC ^min^ Ire1β^+/+^* control mice. We crossed *Lgr5*-eGFP-Cre^ER^-Lox-STOP-Lox-tdtomato mice with *Ire1α*^fl/+^ mice to generate *Lgr5*-eGFP-Cre^ER^-Lox-STOP-Lox-tdtomato-*Ire1α*^fl/fl^ (eGFP-tdtomato-*Ire1α^-/-^*) mice and littermate *Lgr5*-EGFP-Cre^ER^-Lox-STOP-Lox-tdtomato-*Ire1α*^+/+^(eGFP-tdtomato-*Ire1α*^+/+^) mice for experiments. All mice were maintained in specific pathogen-free rooms at the Sanford Burnham Prebys Medical Discovery Institute animal facility. When we harvested tissue, mice were euthanized by CO_2_ inhalation. All animal protocols were reviewed and approved by the Institutional Animal Care and Use Committee at the SBP Medical Discovery Institute.

### Histology staining

Formalin-fixed and paraffin embedded sections were prepared by the histology core facility at the SBP Medical Discovery Institute. These paraffin sections were used for hematoxylin and eosin (H&E) and Alcian Blue staining. All staining was performed by experienced personnel of the same core facility. Images were taken by using Nikon TE300 and Olympus IX81 microscopes in the imaging core facility at SBP.

### Immunofluorescence staining

Formalin fixed paraffin embedded sections were used for immunofluorescence staining. Sections were dewaxed and rehydrated first and incubated with primary antibodies at 4°C overnight. After washing carefully with PBS, samples were incubated at room temperature for 2 hours with goat anti rabbit (Cat # A11012, Invitrogen or Cat# DI-1488 Vector lab) or goat anti rat (SA5-10020 Invitrogen) secondary antibodies. Images were taken using a Nikon TE300 microscope at imaging core facility of SBP. Primary antibodies used for immunofluorescence staining were: Ki67 (Cat #12202s, Cell Signaling Technology, USA); LYZ (Cat# A0099 Dako, USA); Ep-CAM (Cat #:118201, Biolegend, USA).

### Fluorescence In Situ Hybridization (FISH)

We collaborated with ADVANCED CELL DIAGNOSTICS, Inc. to design the probes to detect *Ire1α* (ERN1) and *Ire1β* (ERN2). The *Ire1α* probe is a 20 ZZ probe pair that targets region 162 – 1242 of NM_023913.2 that detects part of exon 1 through part of exon 11 which covers ∼27% of the total mature RNA sequence. *Ire1β* is a 20 ZZ probe pair with target region 2 – 958 of NM_001316689.1 that detects part of exon 1 through part of exon 9 which covers ∼34% of the total mature RNA sequence. We collaborated with the Histology Core at Sanford Burnham Prebys Medical Discovery Institute to perform the formalin-fixed paraffin-embedded (FFPE) tissue processing and sectioning prior to FISH, which was conducted following the manufacture’s FISH protocol (# 323100, ACD, USA). *Ire1α* and *Ire1β* probes were applied on the same slide and assigned to different detection channels to identify the expression pattern of these two molecules on the same tissue section. The *Olfm4* probe (# 311831-C2, ACD, USA) was applied on the same slide as the *Ire1α* and *Ire1β* probes to test whether ISCs express these two molecules.

Antibodies for LYZ (Cat# A0099 Dako, USA), MUC2(#NBP1-31231, Novus Biologicals, USA) and Ki67(Cat #12202s, Cell Signaling Technology, USA) were applied on the same slides as *Ire1α* and *Ire1β* probes, each with its respective detection channel. Immunofluorescence staining for LYZ, MUC2 and Ki67 was performed after completing the FISH procedure for *Ire1α* and *Ire1β* probes.

### Transmission electron microscopy

Small intestines and colons were opened longitudinally and cleaned quickly on ice using ice-cold PBS (CORNING, #21-040-CV). Then 3-5mm intestinal segments were treated with fixative obtained from the University of California, San Diego (UCSD) electron microscopy core facility (La Jolla, CA). Tissue segments were sent to the same core for subsequent processing steps by a TEM expert. Images were taken on an electron microscope (JEOL 1400 plus) in the same core facility.

### Isolation of small intestinal epithelial cells

The whole process was modified as previously described(37). In summary, after carefully removing Peyer’s Patches and intestinal contents, the small intestines were opened longitudinally and cut into 2–4 mm pieces. Intestinal segments were washed well in ice-cold PBS and rocked for 30min at 4°C in isolation buffer (2mM EDTA in PBS). After all fragments settled down the supernatant was removed. Then, 20ml of ice-cold PBS was added and pipetted up and down to fully release the villi and crypts into the supernatant. The supernatant was analyzed by Leica inverted microscopy (MS5, Leica, Germany) to ensure they were enriched with villi and crypts. The supernatant was then passed through a 100 μm cell strainer (Celltreat Scientific Products, USA, #229485) and cells were collected into a 1% BSA coated 50mL tube.

### Intestinal permeabilization assay

The intestinal permeability was measured using FITC-labeled dextran. In brief, after food and water were withdrawn for 4h, mice were orally administrated with FITC-labeled dextran 600 mg/kg (# Sigma-Aldrich). Blood was collected 4h after dextran administration and serum FITC-dextran concentrations were measured using a microplate reader (Spectra ID3).

### Measurement of ISC survival advantage

The survival potential of ISCs in crypts from Lgr5-eGFP-Cre^ER^-Lox-STOP-Lox-tdtomato-*IRE1α^fl/fl^* (eGFP-tdtomato-*Ire1α^-/-^*) and Lgr5-EGFP-Cre^ER^-Lox-STOP-Lox-tdtomato-*IRE1α*^+/+^ (eGFP-tdtomato-*Ire1α^+/+^*) mice was measured as described(38). Briefly, mice were injected (IP) with 0.15mg Tamoxifen dissolved in corn oil. Intestines were harvested 20d later and fixed in 4% paraformaldehyde for 4h at RT. Then 1×1 cm fragments were mounted on slides and incubated in PBS containing DAPI 10µg/ml. Intestinal crypt bottoms were visualized on an Olympus confocal microscope (Fluoview 1000) in the imaging core facility at SBP and clone sizes were quantified.

### AOM/DSS model

The AOM/DSS model was used according to previous studies(13). Briefly, 6-8wk-old mice were challenged with 12.5mg/kg Azoxymethane (AOM) (Cat # A5486-25MG, Sigma) through i.p. injection followed by two 7d cycles of 2.5% Dextran Sulfate Sodium Salt (DSS) (Cat#14489, Affymetrix) in drinking water with an interval of 14d for *Ire1α*^-/-^ mice or followed by 6 days of one cycle 2% DSS after AOM injection for *Ire1β^-/-^* mice. Colorectal tumor numbers and areas were measured at 4months after AOM injection.

For short term intestinal stem cell regeneration analysis in *Ire1β^-/-^* mice, 12.5mg/kg AOM was injected into mice and followed by 6 days of 2% DSS treatment. Colons were harvested 21 days after AOM injection and analyzed.

### RT-PCR for *Xbp1* splicing

RNA was isolated using RNeasy Mini Kit (#74104, Qiagen, USA) and 1µg was used for cDNA synthesis. cDNA was synthesized using iScript Reverse Transcription Supermix (#170-8891, Bio-Red Laboratories, USA) and was used for *Xbp1s* PCR analysis. The primers used for the *Xbp1* splicing were, forward: ACACGCTTGGGAATGGACAC, reverse: ACACGCTTGGGAATGGACAC.

### Real-time quantitative RT-PCR

RNA was isolated with the RNeasy Mini Kit (#74104, Qiagen, USA), and 1 µg was used to synthesize cDNA. The iScript Reverse Transcription Supermix (#170-8891, Bio-Rad Laboratories, USA) was used for cDNA synthesis, which was then applied for subsequent quantitative PCR using SYBR Green Supermix (#1725124 Bio-Red Laboratories, USA). Primers used were listed in Table1.

### RNA Sequencing and RNA-Seq data analysis

RNA was extracted from IECs using the RNeasy Mini Kit (#74104, Qiagen, USA) and was sent to AZENTA (USA) for RNA sequencing. RNA-Seq data analysis was conducted through collaboration with the Bioinformatic Core at Sanford Burnham Prebys Medical Discovery Institute. Illumina Truseq adapter, polyA, and polyT sequences were trimmed with cutadapt v2.3 using parameters “cutadapt -j 4 -m 20 --interleaved -a AGATCGGAAGAGCACACGTCTGAACTCCAGTCAC-A AGATCGGAAGAGCGTCGTGTAGGGAAAGAGTGT Fastq1 Fastq2 | cutadapt --interleaved -j 4 -m 20 -a “A(100)” -A “A(100)” - | cutadapt -j 4 -m 20 -a “T(100)” -A “T(100)” -”(39). Trimmed reads were aligned to mouse genome version 38 (mm10) using STAR aligner v2.7.0d_0221 with parameters according to ENCODE long RNA-Seq pipeline (40) (https://github.com/ENCODE-DCC/long-rna-seq-pipeline). Gene expression levels were quantified using RSEM v1.3.1(41). Ensembl gene annotations version 84 was used for alignment and quantification steps. RNA-Seq sequence, alignment, and quantification quality was assessed using FastQC v0.11.5 (https://www.bioinformatics.babraham.ac.uk/projects/fastqc/) and MultiQC v1.8(42). Biological replicate concordance was assessed using principal component analysis (PCA) and pair-wise Pearson correlation analysis. Lowly expressed genes were filtered out by applying the following criterion: estimated counts (from RSEM) ≥ number of samples 5. Filtered estimated read counts from RSEM were compared using the R Bioconductor package DESeq2 v1.22.2 based on generalized linear model and negative binomial distribution(43). Genes with Benjamini-Hochberg corrected p-value < 0.05 and fold change ≥ 2.0 or ≤ 2.0 were selected as differentially expressed genes. The differentially expressed genes were further analyzed using Gene Set Enrichment Analysis (GSEA) C6 oncogenic signature gene sets to evaluate the oncogenic pathways that expand in *Ire1β-*deleted IECs. We employed stringent criteria for the selection of those pathways, with P value<0.05 and FDR (False Discovery Rate)<10%. The Read-split-Run analysis, which uses RNA-Seq data to detect *Xbp1* mRNA splicing, was initially developed through collaboration between our lab and bioinformatic specialists and was previously described(21).

### Statistical analysis

Statistical analysis was performed using GraphPad Prism 9. The unpaired two-tailed Student t test was used to compare means of two groups. Analysis of variance (ANOVA) with Dunnett’s test was used to compare means of multiple groups against a control group. Fisher’s exact test was used to compare the incidence of bleeding feces between *Ire1β^+/+^* and *Ire1β^-/-^* mice with DSS treatment. Log-rank tests were used for the patient survival probability comparison between either *IRE1α*^high^ and *IRE1α*^low^ or *IRE1β*^high^ and *IRE1β*^low^. All p values are clarified in relevant figure legends. P<0.05 was considered a significant difference.

## Acknowledgements

We thank members of the Kaufman lab for constructive insights during this work. We appreciate the outstanding support from the following SBP Core Facilities: Animal Facility, Histology, Bioinformatics, Next Generation Sequencing, and the University of California, San Diego (UCSD) electron microscopy core facility.

## Supplemental Figures

**Fig. S1.**
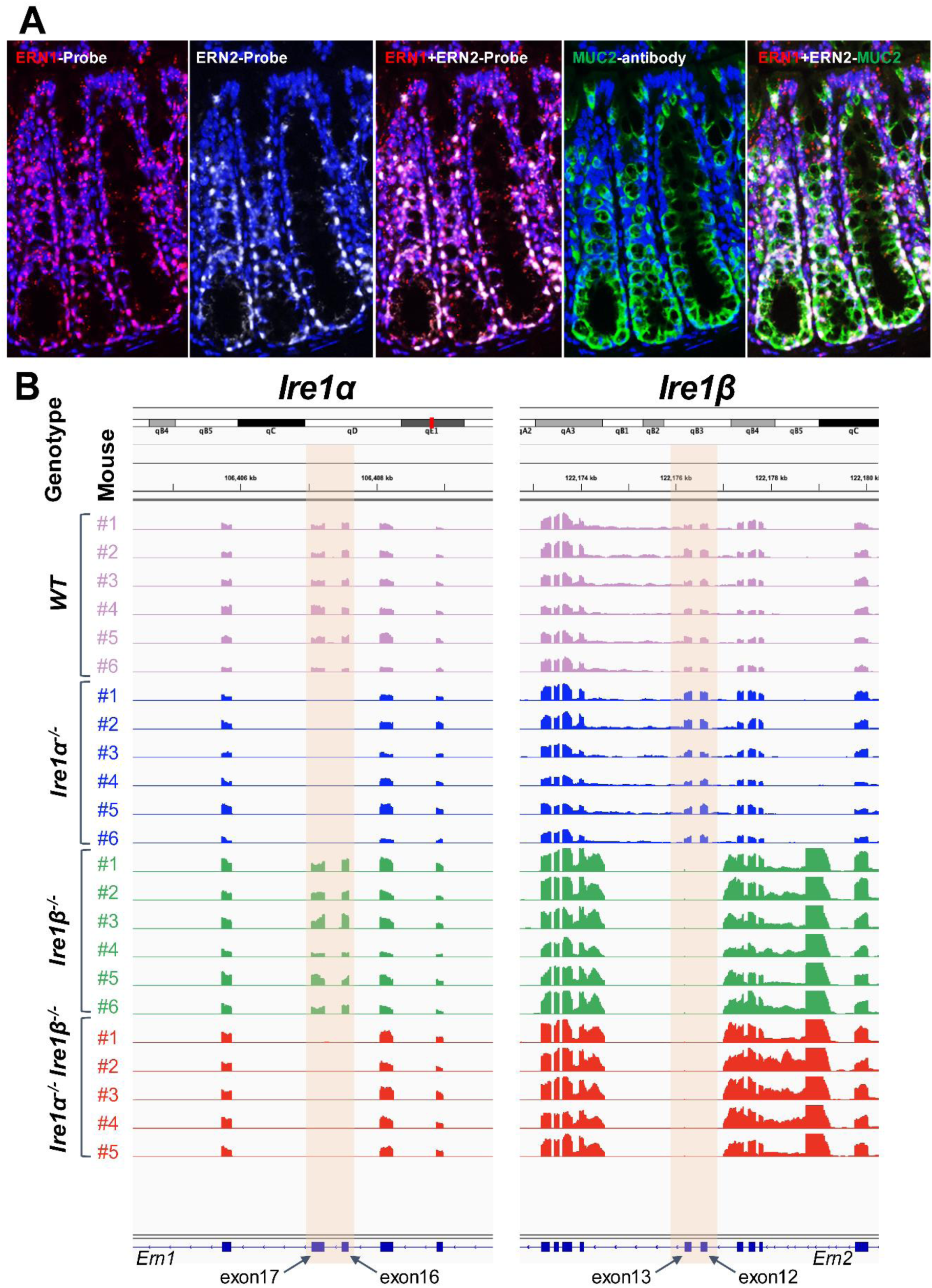
*Ire1α* and *Ire1β* are co-expressed in colon goblet cells and genome browser of RNA-Seq data. (A)The colons of 12-week-old mice were used for FISH to visualize the distribution of *Ire1α* and *Ire1β*. mRNA probes for *Ire1α* and *Ire1β* were applied to the same slide, along with antibody for MUC2, to confirm the expression of two molecules in colon goblet cells. (B) Genome browser showing RNA-Seq coverage at Ire1α and Ire1β loci confirms deletion of exons 16 and 17 in *Ire1α* and exons 12 and 13 in *Ire1β* mice.

**Fig. S2.**
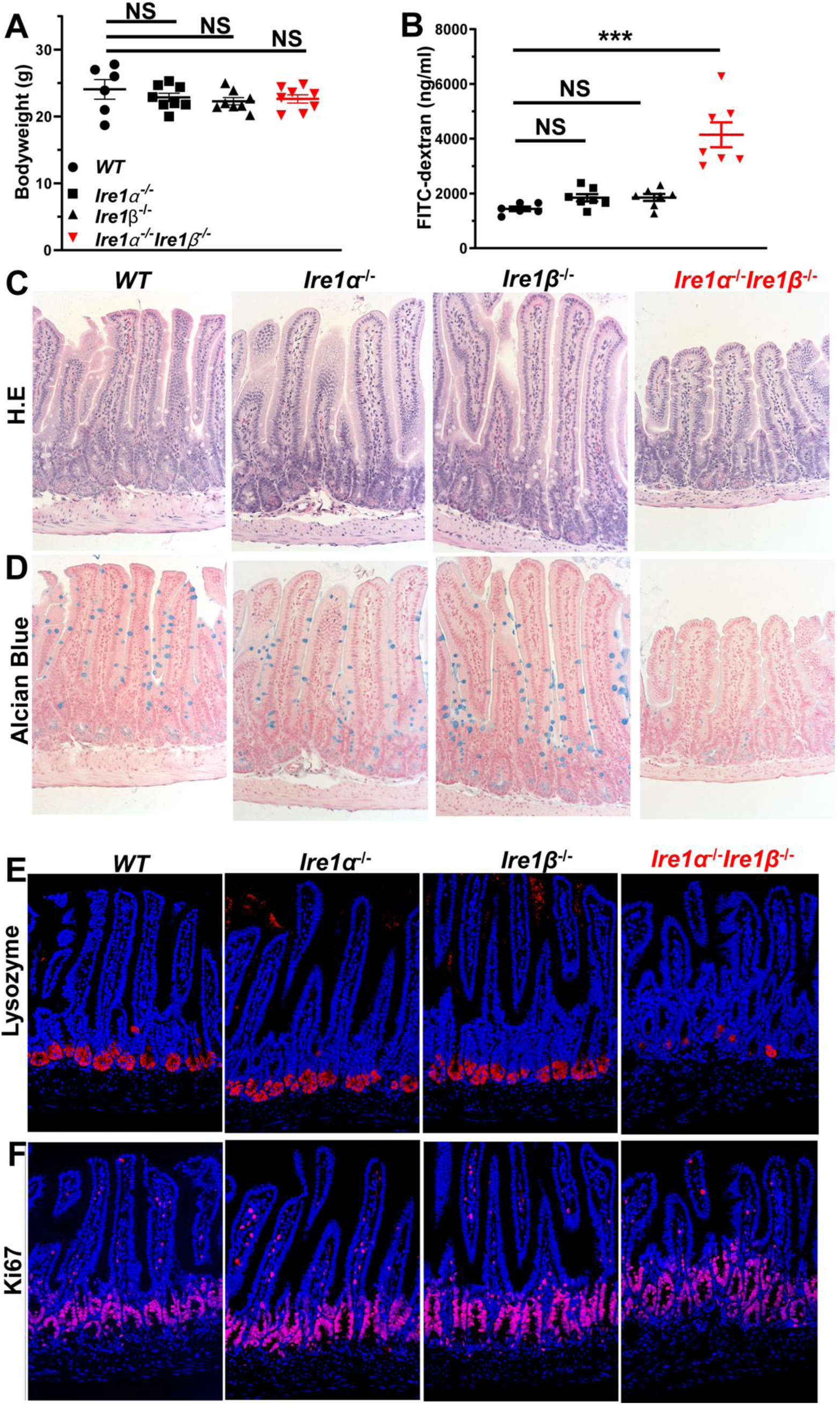
Young adult *Ire1α^-/-^Ire1β^-/-^* mice exhibit small intestinal destruction. (A) Body weight of mice 10-14 wks old, *WT* n=6 mice, *Ire1α^-/-^* n=8 mice, and *Ireβ^-/-^* n= 8 mice, and *Ire1α^-/-^ Ire1β^-/-^* n=8 mice. Data are shown as means ± SE (B) FITC-dextran serum levels (10-14 wks old, *Ire1α^+/+^Ireβ^+/+^* n=7 mice, *Ire1α^-/-^Ireβ^+/+^*, n=7 mice, *Ire1α^+/+^Ireβ^-/-^* n=7 mice, and *Ire1α^-/-^ Ire1β^-/-^* n=7 mice). Data are shown as means ± SE. Serum was collected 4 hours after oral administration of FITC-dextran at 600mg/kg. (C) Small intestine H&E staining, (D) Alcian blue staining, (E) Immunofluorescence of LYZ, (F) Immunofluorescence of Ki67.

**Fig. S3.**
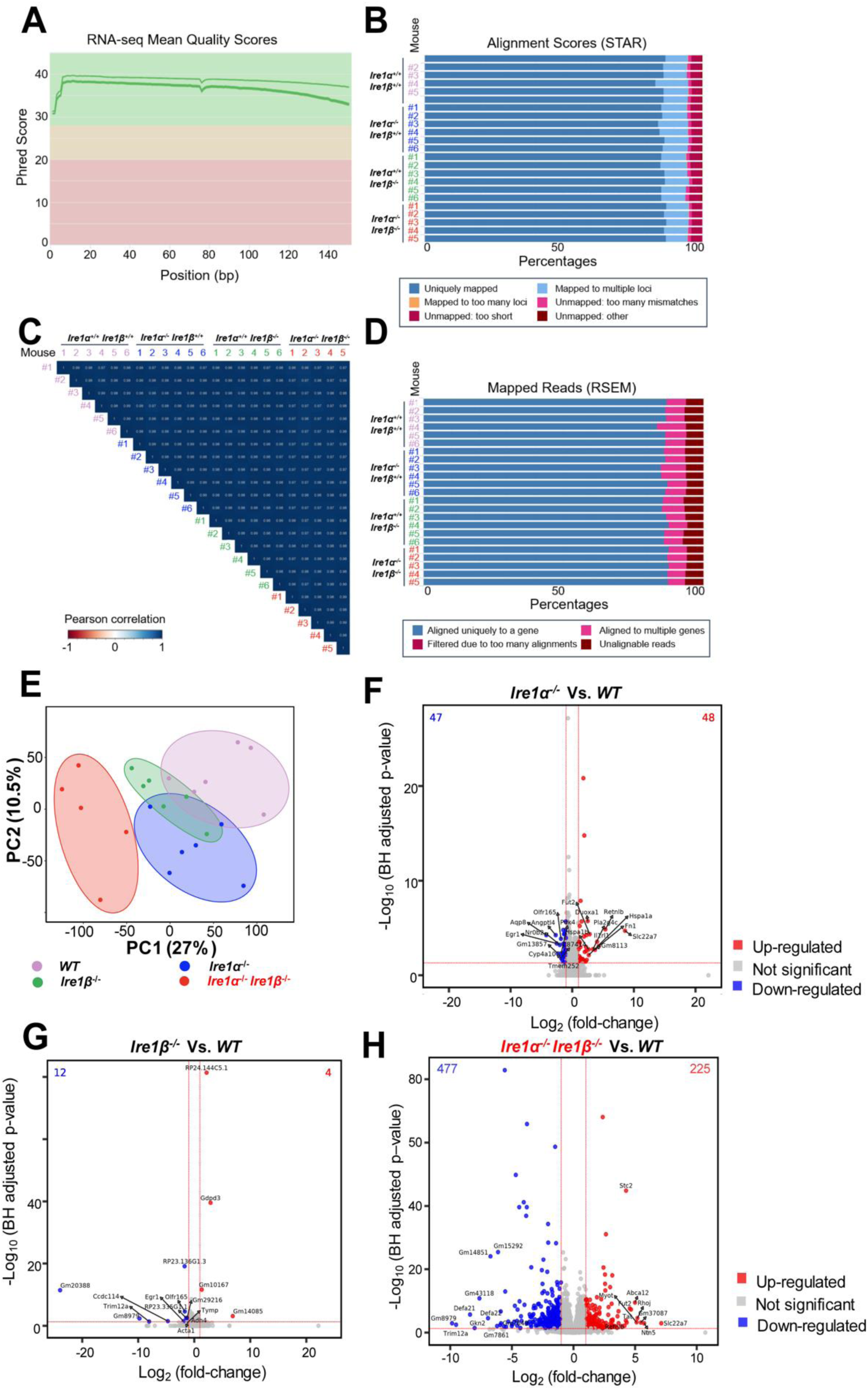
RNA-Seq data without Tm treatment demonstrates quality control. (A) Sequence quality histograms depict the mean quality score at each base pair of the RNA-Seq Illumina reads. (B) Pairwise Pearson correlation coefficients are computed for all samples using transcripts per million (TPM) to depict biological replicate concordance. (C) Breakdown of alignment rates to mouse genome version is shown for all samples using STAR aligner. (D) Breakdown of how reads were mapped to mouse transcriptome version 84 from Ensembl using RSEM is shown. (E) Principal component analysis (PCA) plot shows separation of *Ire1α^-/-^Ireβ^-/-^* from *WT*, *Ire1α^-/-^,* and *Ireβ^-/-^* groups along the main principal component 1 (PC1). Red dots, *Ire1α^-/-^Ireβ^-/-^*; blue dots, *Ire1α^-/-^;* green dots*, Ireβ^-/-^*; purple dots, *WT*. (F) Volcano plots depict –Log_10_ of BH adjusted p-values (y-axis) and Log_2_ of fold-change for *Ire1α^-/-^* versus *WT* comparison. Red and blue dots indicate up-regulated and down-regulated genes respectively using cutoffs of adjusted p-value < 0.05 and abs (Log_2_ fold-change) ≥ 1.0. Gray dots indicate genes that are not significantly changed. The number of up-regulated and down-regulated genes are indicated in the upper right and left corners, respectively. (G) Volcano plot depicts –Log_10_ of BH adjusted p-values (y-axis) and Log_2_ of fold-change for *Ire1β^-/-^* versus *WT* comparison. The legend and plot information are the same as in “E”. (H) Volcano plot summarizes differential gene expression (DE) results of *Ire1α^-/-^Ireβ^-/-^* versus *WT*. Genes with absolute (fold-change) ≥ 2.0 and BH-adjusted p-value < 0.05 were considered as DE.

**Fig. S4.**
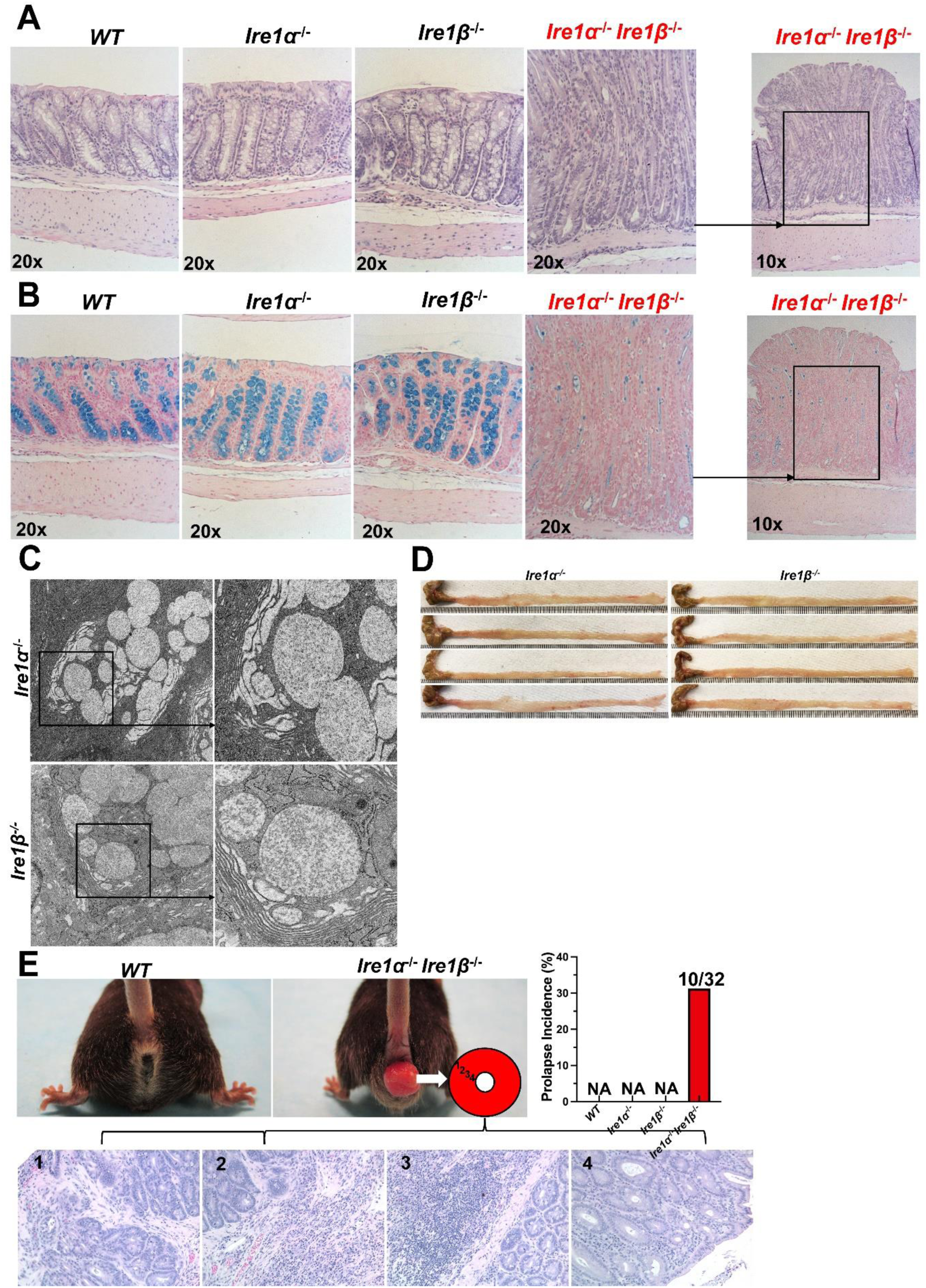
*Ire1α^-/-^ Ire1β^-/-^* mice display extensive colon destruction (A) Colons of young adult mice (10-14 wks old) were harvested for (A) H&E staining and (B) Alcian blue staining. (C)TEM images show secretory particles are well preserved in *Ire1α^-/-^* and *Ire1β^-/-^* single deleted mice. Colons of 14 wks old mice were used for TEM analysis (n=2 mice). (D) Representative colon images of mice older than 40 wks old are shown. (E) Spontaneous prolapse occurs in *Ire1α^-/-^ Ireβ^-/-^* mice. Mouse images show prolapse in *Ire1α^-/-^Ireβ^-/-^* mice, the incidence is about 40% (10 out of 32 mice). H&E staining shows inflammation (2,3) and residual intestinal gland of colon (1,4) in prolapse.

**Fig. S5.**
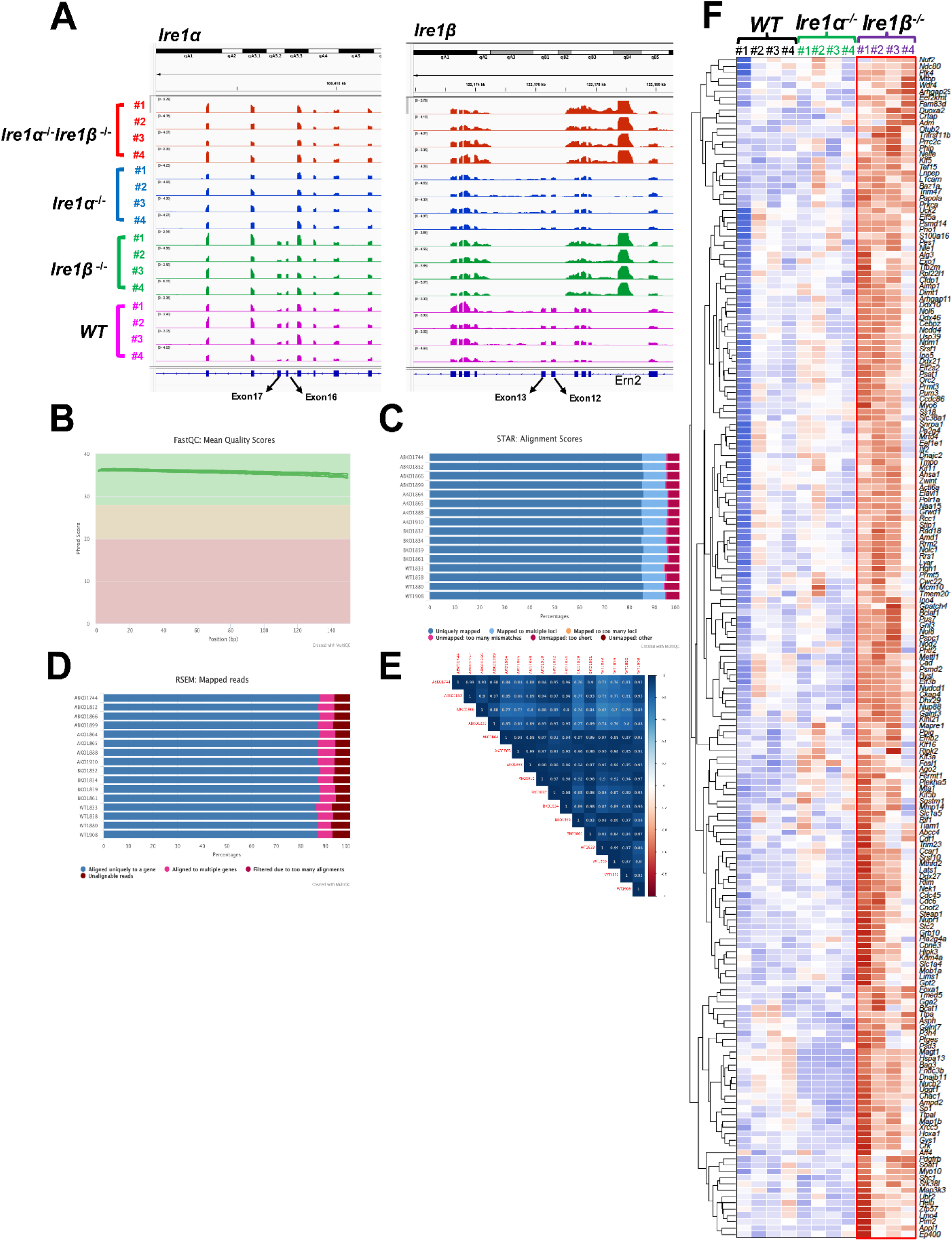
RNA-Seq data is high quality for analysis for Tm-treated IECs mRNA and heatmap of selected genes. (A) Genome browser shows RNA-Seq coverage at *Ire1α* and *Ire1β* loci that confirms deletion of exons 16 and 17 in *Ire1α* and exons 12 and 13 in *Ire1β* mice. (B) Sequence quality histograms depict the mean quality score at each base pair of the RNA-Seq Illumina reads. (C) Pairwise Pearson correlation coefficients were computed for all samples using transcripts per million (TPM) to depict biological replicate concordance. (D) STAR aligner was used to breakdown alignment rates to the mouse genome version mm10 for all samples. (E) Breakdown of how reads were mapped to mouse transcriptome version 84 from Ensembl using RSEM. (F) Heatmap of selected genes from Tm-treated IECs RNA-Seq data that show expansion of the oncogenic gene set in *Ire1β*^-/-^ IECs is a Global change. (n=4 mice for each group).

---

**Table 1.**
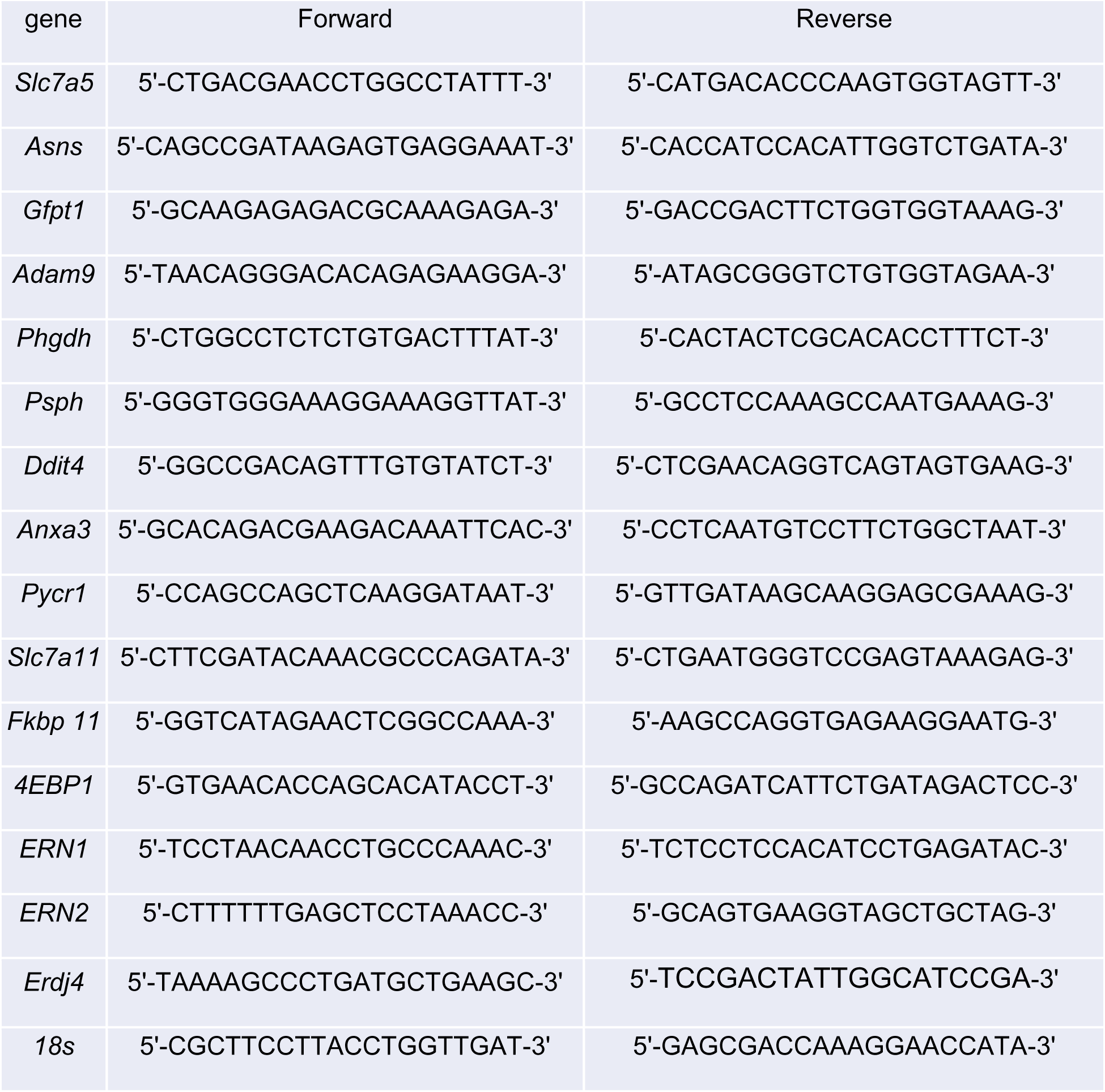
The list of primers used for qPCR.

